## Supplementary material for "Metabolic disruption impairs ribosomal protein levels, resulting in enhanced aminoglycoside tolerance": MIC of antibiotics (from another version of this preprint: version 4, 2024-06-07)

**Supplementary file 1. MIC of antibiotics.**

**Table 1. The minimum inhibitory concentration (MIC) levels of streptomycin, gentamicin, and amikacin were examined in TCA and ETC mutants as well as the wild type.**

| **Strains of *E. coli* MG1655** | **MIC (μg/ml) of  Streptomycin** | **MIC (μg/ml) of  Gentamicin** | **MIC (μg/ml) of  Amikacin** |
| --- | --- | --- | --- |
| Wild type | 2-3 | 0.125-0.19 | 0.75 |
| Δ*sucA* | 2-3 | 0.5-0.75 | 1.5-2 |
| Δ*gltA* | 4 | 0.5-0.75 | 1.5-2 |
| Δ*nuoI* | 6 | 1-1.5 | 3 |
| Δ*nuoM* | 6 | 0.38-0.50 | 2 |
| Δ*mdh* | 3-4 | 0.38-0.50 | 1.5-2 |
| Δ*sdhC* | 3 | 0.38-0.50 | 2-3 |
| Δ*icd* | 3 | 0.15 | 2 |
| Δ*acnB* | 2-3 | 0.25-0.38 | 1.5 |
| Δ*fumA* | 6 | 0.19 | 1-1.5 |
