## Supplementary material for "Metabolic disruption impairs ribosomal protein levels, resulting in enhanced aminoglycoside tolerance": Tables associated with proteomics data (from another version of this preprint: version 4, 2024-06-07)

**Supplementary file 2. Proteomics data.**

**Table 1. Upregulated proteins at mid-exponential phase (t=3.5 h) in the Δ*sucA* mutant strain relative to the wild type.** FC: Fold Change. A significance threshold of P < 0.05, based on F- and t-statistics (see Materials and Methods), is applied.

| **Protein ID** | **Description** | **Log_2_(FC)** |
| --- | --- | --- |
| P21151 | 3-ketoacyl-CoA thiolase FadA | 4.62142 |
| P77581 | Succinylornithine transaminase | 4.08394 |
| P75818 | Uncharacterized lipoprotein YbjP | 3.72890 |
| P07003 | Pyruvate dehydrogenase [ubiquinone] | 3.70453 |
| P25526 | Succinate-semialdehyde dehydrogenase [NADP(+)] GabD | 3.62509 |
| P21367 | Probable hydrolase YcaC | 3.31773 |
| P22256 | 4-aminobutyrate aminotransferase GabT | 3.27743 |
| P75694 | Uncharacterized protein YahO | 3.09843 |
| P21177 | Fatty acid oxidation complex subunit alpha | 2.95728 |
| P61320 | Outer-membrane lipoprotein LolB | 2.95618 |
| P39325 | Galactofuranose-binding protein YtfQ | 2.90126 |
| P0AFM4 | Phosphate starvation-inducible protein PsiF | 2.78265 |
| P0A867 | Transaldolase A | 2.69000 |
| P37330 | Malate synthase G | 2.34830 |
| P76193 | Probable L,D-transpeptidase YnhG | 2.28483 |
| P0A8G6 | NAD(P)H dehydrogenase (quinone) | 2.27339 |
| P75691 | Aldehyde reductase YahK | 2.15509 |
| P39371 | N-acetylneuraminate epimerase | 2.11523 |
| P77717 | Uncharacterized lipoprotein YbaY | 2.11415 |
| P0A9S3 | Galactitol 1-phosphate 5-dehydrogenase | 2.09646 |
| P0ADU5 | Protein YgiW | 2.07362 |
| P0ADB7 | Entericidin B | 2.06776 |
| P0AG80 | sn-glycerol-3-phosphate-binding periplasmic protein UgpB | 2.05784 |
| P08506 | D-alanyl-D-alanine carboxypeptidase DacC | 2.02760 |
| P0ADE6 | Potassium binding protein Kbp | 2.01108 |
| P0AFH8 | Osmotically-inducible protein Y | 1.97533 |
| P0ADI4 | Enterobactin synthase component B | 1.96131 |
| P25906 | Pyridoxine 4-dehydrogenase | 1.94299 |
| P0ADX7 | Uncharacterized protein YhhA | 1.94256 |
| P0A991 | Fructose-bisphosphate aldolase class 1 | 1.94223 |
| P0A9S1 | Lactaldehyde reductase | 1.86026 |
| P10908 | Glycerophosphodiester phosphodiesterase, cytoplasmic | 1.77529 |
| P23524 | Glycerate 2-kinase | 1.77020 |
| P27550 | Acetyl-coenzyme A synthetase | 1.76624 |
| P23522 | 5-keto-4-deoxy-D-glucarate aldolase | 1.69987 |
| P0AES2 | Glucarate dehydratase | 1.67747 |
| P23847 | Dipeptide-binding protein | 1.59291 |
| P0C0L2 | Peroxiredoxin OsmC | 1.58904 |
| P02943 | Maltoporin | 1.47922 |
| P42620 | Glutathionyl-hydroquinone reductase YqjG | 1.46141 |
| P31658 | Protein/nucleic acid deglycase 1 | 1.42126 |
| P0C058 | Small heat shock protein IbpB | 1.40640 |
| P0ABT2 | DNA protection during starvation protein | 1.39667 |
| P0AB24 | Iron uptake system component EfeO | 1.38413 |
| P0A6J5 | D-amino acid dehydrogenase | 1.36382 |
| P37188 | PTS system galactitol-specific EIIB component | 1.35253 |
| P0ABQ2 | 2-hydroxy-3-oxopropionate reductase | 1.35170 |
| P69828 | PTS system galactitol-specific EIIA component | 1.27256 |
| P09551 | Lysine/arginine/ornithine-binding periplasmic protein | 1.26676 |
| P25516 | Aconitate hydratase A | 1.25936 |
| P0A879 | Tryptophan synthase beta chain | 1.22606 |
| P68206 | UPF0337 protein YjbJ | 1.22500 |
| P19926 | Glucose-1-phosphatase | 1.22424 |
| P0C8J6 | D-tagatose-1,6-bisphosphate aldolase subunit GatY | 1.20844 |
| P0A9G6 | Isocitrate lyase | 1.17487 |
| P08997 | Malate synthase A | 1.16133 |
| P07014 | Succinate dehydrogenase iron-sulfur subunit | 1.10148 |
| P00370 | NADP-specific glutamate dehydrogenase | 1.09470 |
| P25553 | Lactaldehyde dehydrogenase | 1.08900 |
| P32162 | UPF0381 protein YiiS | 1.08540 |
| P29012 | Alanine racemase, catabolic | 1.04998 |
| P0AC41 | Succinate dehydrogenase flavoprotein subunit | 1.00542 |
| P0AD59 | Inhibitor of vertebrate lysozyme | 1.00289 |
| P0C8J8 | D-tagatose-1,6-bisphosphate aldolase subunit GatZ | 0.99139 |
| Q46845 | Disulfide-bond oxidoreductase YghU | 0.98556 |
| P0A6D3 | 3-phosphoshikimate 1-carboxyvinyltransferase | 0.97868 |
| P0A744 | Peptide methionine sulfoxide reductase MsrA | 0.96690 |
| P77804 | Protein YdgA | 0.94800 |
| P09832 | Glutamate synthase [NADPH] small chain | 0.90847 |
| P05791 | Dihydroxy-acid dehydratase | 0.90739 |
| P69797 | PTS system mannose-specific EIIAB component | 0.87185 |
| P0AB80 | Branched-chain-amino-acid aminotransferase | 0.86286 |
| P0A9Q7 | Bifunctional aldehyde-alcohol dehydrogenase AdhE | 0.79941 |
| P0AC84 | Hydroxyacylglutathione hydrolase GloB | 0.78259 |
| P0ABD3 | Bacterioferritin | 0.76902 |
| P75797 | Glutathione-binding protein GsiB | 0.75115 |
| P76116 | Uncharacterized protein YncE | 0.74836 |
| P31068 | Flagellar assembly protein FliH | 0.73115 |
| P76187 | Oxidoreductase YdhF | 0.72639 |
| P0AFX4 | Regulator of sigma D | 0.72469 |
| P05042 | Fumarate hydratase class II | 0.72092 |
| P0A8W2 | Transcriptional regulator SlyA | 0.71564 |
| P37902 | Glutamate/aspartate import solute-binding protein | 0.71115 |
| P25894 | Metalloprotease LoiP | 0.70036 |
| P06149 | Quinone-dependent D-lactate dehydrogenase | 0.70015 |
| P0A761 | Putative N-acetylmannosamine-6-phosphate 2-epimerase | 0.69313 |
| P23721 | Phosphoserine aminotransferase | 0.69212 |
| P0AEZ9 | Molybdenum cofactor biosynthesis protein B | 0.66728 |
| P39099 | Periplasmic pH-dependent serine endoprotease DegQ | 0.66398 |
| P39315 | Quinone oxidoreductase 2 | 0.66360 |
| P0A8C1 | UPF0145 protein YbjQ | 0.66318 |
| P30859 | Putative ABC transporter arginine-binding protein 2 | 0.65763 |
| P36683 | Aconitate hydratase B | 0.65211 |
| P46853 | Uncharacterized oxidoreductase YhhX | 0.56771 |
| P0A759 | Glucosamine-6-phosphate deaminase | 0.54391 |
| P62707 | 2,3-bisphosphoglycerate-dependent phosphoglycerate mutase | 0.53517 |
| P52697 | 6-phosphogluconolactonase | 0.53211 |
| P06610 | Thioredoxin/glutathione peroxidase BtuE | 0.52276 |
| P0A9Q9 | Aspartate-semialdehyde dehydrogenase | 0.51405 |
| P75960 | NAD-dependent protein deacylase | 0.49760 |
| P18843 | NH(3)-dependent NAD(+) synthetase | 0.49214 |
| P0AC33 | Fumarate hydratase class I, aerobic | 0.49130 |
| P76346 | Protein MtfA | 0.48105 |
| P12758 | Uridine phosphorylase | 0.47557 |
| P0AB91 | Phospho-2-dehydro-3-deoxyheptonate aldolase, Phe-sensitive | 0.45776 |
| P22188 | UDP-N-acetylmuramoyl-L-alanyl-D-glutamate--2,6-diaminopimelate ligase | 0.45749 |
| P45578 | S-ribosylhomocysteine lyase | 0.45466 |
| P23843 | Periplasmic oligopeptide-binding protein | 0.45382 |
| P0AF18 | N-acetylglucosamine-6-phosphate deacetylase | 0.45120 |
| P0AGG4 | Thioredoxin 2 | 0.44531 |
| P0A6K6 | Phosphopentomutase | 0.43619 |
| P0A9L8 | Pyrroline-5-carboxylate reductase | 0.41103 |
| P29745 | Peptidase T | 0.39708 |
| P0ADE8 | tRNA-modifying protein YgfZ | 0.39692 |
| P07024 | Protein UshA | 0.39353 |
| P0AD49 | Ribosome-associated inhibitor A | 0.39338 |
| P0A6S0 | Flagellar L-ring protein | 0.37552 |
| P0AEU7 | Chaperone protein Skp | 0.37464 |
| P0A6F3 | Glycerol kinase | 0.37384 |
| P08331 | 2',3'-cyclic-nucleotide 2'-phosphodiesterase/3'-nucleotidase | 0.36371 |
| P0AEE5 | D-galactose/methyl-galactoside binding periplasmic protein MglB | 0.36347 |
| P07650 | Thymidine phosphorylase | 0.34584 |
| P0A6T1 | Glucose-6-phosphate isomerase | 0.34322 |
| P02925 | Ribose import binding protein RbsB | 0.34155 |
| P27248 | Aminomethyltransferase | 0.33278 |
| P27306 | Soluble pyridine nucleotide transhydrogenase | 0.31942 |
| P0ABF6 | Cytidine deaminase | 0.31797 |
| P00509 | Aspartate aminotransferase | 0.28581 |
| P0AE08 | Alkyl hydroperoxide reductase C | 0.27486 |
| P75874 | Uncharacterized protein YccU | 0.20902 |

**Table 2. Upregulated proteins at mid-exponential phase (t=3.5 h) in the Δ*gltA* mutant strain relative to the wild type.** FC: Fold Change. A significance threshold of P < 0.05 is applied.

| **Protein ID** | **Description** | **Log_2_(FC)** |
| --- | --- | --- |
| P21151 | 3-ketoacyl-CoA thiolase FadA | 4.60667 |
| P61320 | Outer-membrane lipoprotein LolB | 3.27619 |
| P75818 | Uncharacterized lipoprotein YbjP | 3.16578 |
| P21177 | Fatty acid oxidation complex subunit alpha | 2.83287 |
| P0AFM4 | Phosphate starvation-inducible protein PsiF | 2.29213 |
| P77581 | Succinylornithine transaminase | 2.15198 |
| P25526 | Succinate-semialdehyde dehydrogenase [NADP(+)] GabD | 2.13292 |
| P39325 | Galactofuranose-binding protein YtfQ | 2.03533 |
| P27550 | Acetyl-coenzyme A synthetase | 1.94548 |
| P76193 | Probable L,D-transpeptidase YnhG | 1.89651 |
| P21367 | Probable hydrolase YcaC | 1.89411 |
| P22256 | 4-aminobutyrate aminotransferase GabT | 1.87103 |
| P37330 | Malate synthase G | 1.79141 |
| P13445 | RNA polymerase sigma factor RpoS | 1.75738 |
| P06988 | Histidinol dehydrogenase | 1.73918 |
| P0ADU5 | Protein YgiW | 1.66825 |
| P75694 | Uncharacterized protein YahO | 1.60585 |
| P77717 | Uncharacterized lipoprotein YbaY | 1.54546 |
| P0AG80 | sn-glycerol-3-phosphate-binding periplasmic protein UgpB | 1.41793 |
| P0AF03 | Molybdopterin adenylyltransferase | 1.40496 |
| P0ADE6 | Potassium binding protein Kbp | 1.38580 |
| P0A9S1 | Lactaldehyde reductase | 1.37939 |
| P0A9U6 | HTH-type transcriptional regulator PuuR | 1.37286 |
| P0AD96 | Leu/Ile/Val-binding protein | 1.37284 |
| P37902 | Glutamate/aspartate import solute-binding protein | 1.30815 |
| P0AFH8 | Osmotically-inducible protein Y | 1.30741 |
| P0A867 | Transaldolase A | 1.27951 |
| P05042 | Fumarate hydratase class II | 1.27550 |
| P00370 | NADP-specific glutamate dehydrogenase | 1.23009 |
| P76004 | Uncharacterized protein YcgM | 1.18993 |
| P09127 | Protein HemX | 1.15950 |
| P0A6T9 | Glycine cleavage system H protein | 1.15232 |
| P39371 | N-acetylneuraminate epimerase | 1.12458 |
| P0ADZ7 | Sec translocon accessory complex subunit YajC | 1.10859 |
| P23847 | Dipeptide-binding protein | 1.10537 |
| P68206 | UPF0337 protein YjbJ | 1.08328 |
| P68187 | Maltose/maltodextrin import ATP-binding protein MalK | 1.06419 |
| P75691 | Aldehyde reductase YahK | 1.02785 |
| P09551 | Lysine/arginine/ornithine-binding periplasmic protein | 1.01410 |
| P0AC02 | Outer membrane protein assembly factor BamD | 0.97804 |
| P0A9C5 | Glutamine synthetase | 0.96149 |
| P60340 | tRNA pseudouridine synthase B | 0.94336 |
| P0ADB7 | Entericidin B | 0.91997 |
| P23843 | Periplasmic oligopeptide-binding protein | 0.90332 |
| P0ABT2 | DNA protection during starvation protein | 0.89360 |
| P25553 | Lactaldehyde dehydrogenase | 0.84895 |
| P0AAI3 | ATP-dependent zinc metalloprotease FtsH | 0.83839 |
| P29744 | Flagellar hook-associated protein 3 | 0.80092 |
| P61889 | Malate dehydrogenase | 0.78278 |
| P19926 | Glucose-1-phosphatase | 0.77727 |
| P09546 | Bifunctional protein PutA | 0.77537 |
| P0AB24 | Iron uptake system component EfeO | 0.77490 |
| P0AC33 | Fumarate hydratase class I, aerobic | 0.77302 |
| P39286 | Small ribosomal subunit biogenesis GTPase RsgA | 0.77146 |
| P0AFX4 | Regulator of sigma D | 0.75916 |
| P0AEE1 | Inner membrane lipoprotein DcrB | 0.75837 |
| P25516 | Aconitate hydratase A | 0.75810 |
| P0AC41 | Succinate dehydrogenase flavoprotein subunit | 0.73301 |
| P33235 | Flagellar hook-associated protein 1 | 0.72727 |
| P0A6J5 | D-amino acid dehydrogenase | 0.71079 |
| P0AF50 | Uncharacterized protein YjbR | 0.70914 |
| P0A991 | Fructose-bisphosphate aldolase class 1 | 0.70656 |
| P39265 | D-allose-binding periplasmic protein | 0.70500 |
| P0AGE9 | Succinate--CoA ligase [ADP-forming] subunit alpha | 0.69484 |
| P0A879 | Tryptophan synthase beta chain | 0.68852 |
| P0AD59 | Inhibitor of vertebrate lysozyme | 0.68552 |
| P07014 | Succinate dehydrogenase iron-sulfur subunit | 0.67033 |
| P0AEU0 | Histidine-binding periplasmic protein | 0.66854 |
| P0AGG4 | Thioredoxin 2 | 0.66500 |
| P0A9B6 | D-erythrose-4-phosphate dehydrogenase | 0.64740 |
| P77554 | Uncharacterized protein YahJ | 0.64184 |
| P0A8G3 | Uronate isomerase | 0.61314 |
| P10346 | Glutamine transport ATP-binding protein GlnQ | 0.61090 |
| P0A940 | Outer membrane protein assembly factor BamA | 0.60234 |
| P0A8G6 | NAD(P)H dehydrogenase (quinone) | 0.59030 |
| P64483 | Multifunctional Ser/Thr-tRNA deacylase ProXp-y | 0.58990 |
| P25894 | Metalloprotease LoiP | 0.58142 |
| P62768 | UPF0325 protein YaeH | 0.57908 |
| P36683 | Aconitate hydratase B | 0.56923 |
| P29012 | Alanine racemase, catabolic | 0.56476 |
| P64596 | Outer membrane lipoprotein DolP | 0.55619 |
| P04983 | Ribose import ATP-binding protein RbsA | 0.55488 |
| P0ABD3 | Bacterioferritin | 0.54739 |
| P0C0L2 | Peroxiredoxin OsmC | 0.54392 |
| P46130 | Putative acyl-CoA thioester hydrolase YbhC | 0.53373 |
| P0AE22 | Class B acid phosphatase | 0.53180 |
| P76187 | Oxidoreductase YdhF | 0.52918 |
| P76116 | Uncharacterized protein YncE | 0.50631 |
| P61517 | Carbonic anhydrase 2 | 0.49622 |
| P0A836 | Succinate--CoA ligase [ADP-forming] subunit beta | 0.49284 |
| P0A9L8 | Pyrroline-5-carboxylate reductase | 0.48048 |
| P41409 | Pyrimidine-specific ribonucleoside hydrolase RihA | 0.47089 |
| P39160 | D-mannonate oxidoreductase | 0.46351 |
| P0ACB0 | Replicative DNA helicase | 0.46268 |
| P0ABC7 | Modulator of FtsH protease HflK | 0.46123 |
| P0AEE5 | D-galactose/methyl-galactoside binding periplasmic protein MglB | 0.45960 |
| P24232 | Flavohemoprotein | 0.43339 |
| P0AFK0 | Metalloprotease PmbA | 0.42729 |
| P0A752 | Nicotinate-nucleotide adenylyltransferase | 0.41503 |
| P0A746 | Peptide methionine sulfoxide reductase MsrB | 0.40219 |
| P27248 | Aminomethyltransferase | 0.37918 |
| Q46845 | Disulfide-bond oxidoreductase YghU | 0.37651 |
| P30859 | Putative ABC transporter arginine-binding protein 2 | 0.37625 |
| P76346 | Protein MtfA | 0.36474 |
| P0ACC3 | Iron-sulfur cluster insertion protein ErpA | 0.36378 |
| P0A9P0 | Dihydrolipoyl dehydrogenase | 0.35135 |
| P76177 | Protein YdgH | 0.34664 |
| P07017 | Methyl-accepting chemotaxis protein II | 0.34631 |
| P02925 | Ribose import binding protein RbsB | 0.34520 |
| P03841 | Maltose operon periplasmic protein | 0.33115 |
| P0A799 | Phosphoglycerate kinase | 0.32598 |
| P0A968 | Cold shock-like protein CspD | 0.32379 |
| P06149 | Quinone-dependent D-lactate dehydrogenase | 0.31717 |
| P07024 | Protein UshA | 0.30946 |
| P18843 | NH(3)-dependent NAD(+) synthetase | 0.30357 |
| P75823 | Low specificity L-threonine aldolase | 0.30093 |
| P0A6S0 | Flagellar L-ring protein | 0.28805 |
| P38489 | Oxygen-insensitive NAD(P)H nitroreductase | 0.26376 |
| P0AG99 | Protein-export membrane protein SecG | 0.23676 |
| P0A9B2 | Glyceraldehyde-3-phosphate dehydrogenase A | 0.20031 |
| P0A877 | Tryptophan synthase alpha chain | 0.19227 |
| P0AB77 | 2-amino-3-ketobutyrate coenzyme A ligase | 0.19103 |
| Q46868 | Ubiquinone biosynthesis accessory factor UbiK | 0.18435 |
| P0AGD7 | Signal recognition particle protein | 0.14556 |
| P46853 | Uncharacterized oxidoreductase YhhX | 0.13896 |
| P45578 | S-ribosylhomocysteine lyase | 0.13788 |

**Table 3. Upregulated proteins at mid-exponential phase (t=3.5 h) in the Δ*nuoI* mutant strain relative to the wild type.** FC: Fold Change. A significance threshold of P < 0.05 is applied.

| **Protein ID** | **Description** | **Log_2_(FC)** |
| --- | --- | --- |
| P21151 | 3-ketoacyl-CoA thiolase FadA | 4.03749 |
| P75818 | Uncharacterized lipoprotein YbjP | 3.31614 |
| P0ADU5 | Protein YgiW | 2.35399 |
| P0A9S3 | Galactitol 1-phosphate 5-dehydrogenase | 2.25647 |
| P07003 | Pyruvate dehydrogenase [ubiquinone] | 2.10082 |
| P0A6T9 | Glycine cleavage system H protein | 1.90885 |
| P68187 | Maltose/maltodextrin import ATP-binding protein MalK | 1.83239 |
| Q47690 | Homocysteine S-methyltransferase | 1.78372 |
| P0ADE6 | Potassium binding protein Kbp | 1.74168 |
| P0AAT9 | Uncharacterized protein YbeL | 1.67073 |
| P76004 | Uncharacterized protein YcgM | 1.66246 |
| P25748 | HTH-type transcriptional regulator GalS | 1.66183 |
| P15070 | Flagellar motor switch protein FliN | 1.63856 |
| P52613 | Flagellar FliJ protein | 1.63205 |
| P0ADZ7 | Sec translocon accessory complex subunit YajC | 1.59456 |
| P65367 | Flavodoxin YqcA | 1.59212 |
| P45464 | Penicillin-binding protein activator LpoA | 1.57806 |
| P0A9S1 | Lactaldehyde reductase | 1.54898 |
| P0A867 | Transaldolase A | 1.54495 |
| P25397 | Tellurite methyltransferase | 1.52738 |
| P68206 | UPF0337 protein YjbJ | 1.51853 |
| P0AFH8 | Osmotically-inducible protein Y | 1.51434 |
| P22256 | 4-aminobutyrate aminotransferase GabT | 1.49994 |
| P37330 | Malate synthase G | 1.48591 |
| P75938 | Flagellar basal-body rod protein FlgF | 1.47530 |
| P0AC02 | Outer membrane protein assembly factor BamD | 1.47351 |
| P76187 | Oxidoreductase YdhF | 1.45976 |
| P0AD12 | Protein YeeZ | 1.40384 |
| P0A731 | Methylglyoxal synthase | 1.34919 |
| P75691 | Aldehyde reductase YahK | 1.31582 |
| P08506 | D-alanyl-D-alanine carboxypeptidase DacC | 1.30334 |
| P19926 | Glucose-1-phosphatase | 1.29985 |
| P0A972 | Cold shock-like protein CspE | 1.24677 |
| P0AAQ2 | Putative HNH nuclease YajD | 1.24371 |
| P0ADA3 | Murein hydrolase activator NlpD | 1.23046 |
| P30177 | Uncharacterized protein YbiB | 1.22555 |
| P24232 | Flavohemoprotein | 1.22366 |
| P31658 | Protein/nucleic acid deglycase 1 | 1.17959 |
| P0C0L2 | Peroxiredoxin OsmC | 1.17564 |
| P24215 | Mannonate dehydratase | 1.17181 |
| P06988 | Histidinol dehydrogenase | 1.15434 |
| P29012 | Alanine racemase, catabolic | 1.14560 |
| P0C8J8 | D-tagatose-1,6-bisphosphate aldolase subunit GatZ | 1.13966 |
| P0C037 | Pyrimidine/purine nucleoside phosphorylase | 1.13216 |
| P62768 | UPF0325 protein YaeH | 1.12370 |
| P0ACF0 | DNA-binding protein HU-alpha | 1.11287 |
| P0AFM4 | Phosphate starvation-inducible protein PsiF | 1.08056 |
| P0AD49 | Ribosome-associated inhibitor A | 1.05297 |
| P0A799 | Phosphoglycerate kinase | 1.04978 |
| P39265 | D-allose-binding periplasmic protein | 1.03634 |
| P23847 | Dipeptide-binding protein | 1.02473 |
| P18843 | NH(3)-dependent NAD(+) synthetase | 1.01857 |
| P77695 | Protein GnsB | 1.01832 |
| P31068 | Flagellar assembly protein FliH | 1.01799 |
| P0A7R5 | 30S ribosomal protein S10 | 1.00324 |
| P61889 | Malate dehydrogenase | 0.99116 |
| P0AET8 | 7alpha-hydroxysteroid dehydrogenase | 0.97476 |
| P0ADA1 | Thioesterase 1/protease 1/lysophospholipase L1 | 0.95512 |
| P0A7B8 | ATP-dependent protease subunit HslV | 0.94620 |
| P0A7J7 | 50S ribosomal protein L11 | 0.94427 |
| P0A8G6 | NAD(P)H dehydrogenase (quinone) | 0.93970 |
| P0A6L2 | 4-hydroxy-tetrahydrodipicolinate synthase | 0.92874 |
| P64554 | 7-carboxy-7-deazaguanine synthase | 0.90642 |
| P27248 | Aminomethyltransferase | 0.90387 |
| P0A6Q3 | 3-hydroxydecanoyl-[acyl-carrier-protein] dehydratase | 0.89432 |
| P36672 | PTS system trehalose-specific EIIBC component | 0.89405 |
| P52614 | Flagellar hook-length control protein | 0.87098 |
| P0A780 | Transcription antitermination protein NusB | 0.86822 |
| P0AEM4 | Negative regulator of flagellin synthesis | 0.85890 |
| P0AGG8 | Metalloprotease TldD | 0.85180 |
| P0ADU2 | Probable quinol monooxygenase YgiN | 0.85103 |
| P0A8G3 | Uronate isomerase | 0.83413 |
| P33235 | Flagellar hook-associated protein 1 | 0.82789 |
| P0ABD8 | Biotin carboxyl carrier protein of acetyl-CoA carboxylase | 0.82580 |
| P0AF50 | Uncharacterized protein YjbR | 0.82442 |
| P0AG55 | 50S ribosomal protein L6 | 0.81917 |
| P56580 | PTS system glucitol/sorbitol-specific EIIB component | 0.81611 |
| P0A8W8 | UPF0304 protein YfbU | 0.81141 |
| P0A6F3 | Glycerol kinase | 0.80843 |
| P0C0R7 | Ribosomal RNA large subunit methyltransferase E | 0.80754 |
| P69783 | PTS system glucose-specific EIIA component | 0.80472 |
| P07004 | Gamma-glutamyl phosphate reductase | 0.80340 |
| P76177 | Protein YdgH | 0.80283 |
| P0AEZ9 | Molybdenum cofactor biosynthesis protein B | 0.78418 |
| P0A8W2 | Transcriptional regulator SlyA | 0.77167 |
| P0ACC3 | Iron-sulfur cluster insertion protein ErpA | 0.76743 |
| P0A9B2 | Glyceraldehyde-3-phosphate dehydrogenase A | 0.76513 |
| P04983 | Ribose import ATP-binding protein RbsA | 0.76108 |
| P0A7T7 | 30S ribosomal protein S18 | 0.74861 |
| P37440 | Oxidoreductase UcpA | 0.74436 |
| P0A955 | KHG/KDPG aldolase | 0.74410 |
| P0A7G2 | 30S ribosome-binding factor | 0.74092 |
| P0AE52 | Peroxiredoxin Bcp | 0.71760 |
| P0AGE0 | Single-stranded DNA-binding protein | 0.71347 |
| P0A7J3 | 50S ribosomal protein L10 | 0.69275 |
| P75957 | Lipoprotein-releasing system ATP-binding protein LolD | 0.68678 |
| P68066 | Autonomous glycyl radical cofactor | 0.68588 |
| P38489 | Oxygen-insensitive NAD(P)H nitroreductase | 0.68350 |
| P0A752 | Nicotinate-nucleotide adenylyltransferase | 0.67726 |
| P0AGE9 | Succinate--CoA ligase [ADP-forming] subunit alpha | 0.66730 |
| P05706 | PTS system glucitol/sorbitol-specific EIIA component | 0.64961 |
| P29217 | UPF0502 protein YceH | 0.64865 |
| P39160 | D-mannonate oxidoreductase | 0.64476 |
| P0ACJ8 | DNA-binding transcriptional dual regulator CRP | 0.64254 |
| P0A9A9 | Ferric uptake regulation protein | 0.64060 |
| P03841 | Maltose operon periplasmic protein | 0.63985 |
| P0A8A0 | Probable transcriptional regulatory protein YebC | 0.63555 |
| P0A9Z1 | Nitrogen regulatory protein P-II 1 | 0.63104 |
| P25894 | Metalloprotease LoiP | 0.61692 |
| P0AGD7 | Signal recognition particle protein | 0.61668 |
| P0A7R1 | 50S ribosomal protein L9 | 0.61189 |
| P0AE22 | Class B acid phosphatase | 0.60975 |
| P61517 | Carbonic anhydrase 2 | 0.60845 |
| P0ABA6 | ATP synthase gamma chain | 0.60469 |
| P0AEU7 | Chaperone protein Skp | 0.60120 |
| P0AB52 | Protein YchN | 0.59380 |
| P0AD61 | Pyruvate kinase I | 0.58400 |
| P0ABQ4 | Dihydrofolate reductase | 0.57646 |
| P0ADE8 | tRNA-modifying protein YgfZ | 0.57153 |
| P09551 | Lysine/arginine/ornithine-binding periplasmic protein | 0.55055 |
| P77252 | Uncharacterized protein YkgE | 0.53381 |
| P02925 | Ribose import binding protein RbsB | 0.52815 |
| P06983 | Porphobilinogen deaminase | 0.51600 |
| P0A964 | Chemotaxis protein CheW | 0.51076 |
| P46853 | Uncharacterized oxidoreductase YhhX | 0.49162 |
| P24216 | Flagellar hook-associated protein 2 | 0.49083 |
| P0ABQ2 | 2-hydroxy-3-oxopropionate reductase | 0.48913 |
| P09394 | Glycerophosphodiester phosphodiesterase, periplasmic | 0.47493 |
| P00490 | Maltodextrin phosphorylase | 0.47236 |
| P07862 | D-alanine--D-alanine ligase B | 0.45998 |
| P0A9J6 | Ribokinase | 0.45750 |
| P00370 | NADP-specific glutamate dehydrogenase | 0.44616 |
| P63020 | Fe/S biogenesis protein NfuA | 0.44311 |
| P0A877 | Tryptophan synthase alpha chain | 0.44276 |
| P45523 | FKBP-type peptidyl-prolyl cis-trans isomerase FkpA | 0.42775 |
| P75849 | Hydroxyacylglutathione hydrolase GloC | 0.41268 |
| Q46868 | Ubiquinone biosynthesis accessory factor UbiK | 0.40102 |
| P0ADS2 | Cell division protein ZapA | 0.36709 |
| P0AEZ3 | Septum site-determining protein MinD | 0.35187 |
| P0AC41 | Succinate dehydrogenase flavoprotein subunit | 0.34496 |
| P0A9P0 | Dihydrolipoyl dehydrogenase | 0.33907 |
| P0AC33 | Fumarate hydratase class I, aerobic | 0.33063 |
| P0ABK5 | Cysteine synthase A | 0.32972 |
| P0AC62 | Glutaredoxin 3 | 0.32924 |
| P0C0V0 | Periplasmic serine endoprotease DegP | 0.32827 |
| P69503 | Adenine phosphoribosyltransferase | 0.32691 |
| P0ABZ6 | Chaperone SurA | 0.31639 |
| P39831 | NADP-dependent 3-hydroxy acid dehydrogenase YdfG | 0.29828 |
| P0ACD4 | Iron-sulfur cluster assembly scaffold protein IscU | 0.28906 |
| P0ACC7 | Bifunctional protein GlmU | 0.28600 |
| P0A7W7 | 30S ribosomal protein S8 | 0.24851 |
| P09372 | Protein GrpE | 0.21218 |
| P0AE08 | Alkyl hydroperoxide reductase C | 0.19759 |
| P45748 | Threonylcarbamoyl-AMP synthase | 0.19286 |
| P0A9A6 | Cell division protein FtsZ | 0.18833 |

**Table 4. Upregulated proteins at mid-exponential phase (t=3.5 h) in the Δ*icd* mutant strain relative to the wild type.** FC: Fold Change. A significance threshold of P < 0.05 is applied.

| **Protein ID** | **Description** | **Log_2_(FC)** |
| --- | --- | --- |
| P75818 | Uncharacterized lipoprotein YbjP | 2.50234 |
| P37760 | dTDP-4-dehydrorhamnose reductase | 2.04007 |
| P69811 | Multiphosphoryl transfer protein | 1.93583 |
| P29208 | o-succinylbenzoate synthase | 1.53734 |
| P0C054 | Small heat shock protein IbpA | 1.50389 |
| P63386 | Intermembrane phospholipid transport system ATP-binding protein MlaF | 1.49629 |
| P39286 | Small ribosomal subunit biogenesis GTPase RsgA | 1.36686 |
| P0AF03 | Molybdopterin adenylyltransferase | 1.20992 |
| P00954 | Tryptophan--tRNA ligase | 1.15978 |
| P64554 | 7-carboxy-7-deazaguanine synthase | 1.15685 |
| P0AEN1 | NAD(P)H-flavin reductase | 1.15565 |
| P22256 | 4-aminobutyrate aminotransferase GabT | 1.11043 |
| P0AFK0 | Metalloprotease PmbA | 1.09570 |
| P07012 | Peptide chain release factor RF2 | 1.07478 |
| P0C058 | Small heat shock protein IbpB | 1.06872 |
| P10121 | Signal recognition particle receptor FtsY | 1.05463 |
| P06999 | ATP-dependent 6-phosphofructokinase isozyme 2 | 1.05379 |
| P0A780 | Transcription antitermination protein NusB | 1.03248 |
| P0ADE6 | Potassium binding protein Kbp | 1.02456 |
| P63177 | 23S rRNA (guanosine-2'-O-)-methyltransferase RlmB | 0.99690 |
| P30744 | L-serine dehydratase 2 | 0.99205 |
| P61517 | Carbonic anhydrase 2 | 0.97748 |
| P08839 | Phosphoenolpyruvate-protein phosphotransferase | 0.95773 |
| P0A6C1 | Endonuclease 4 | 0.93401 |
| P68187 | Maltose/maltodextrin import ATP-binding protein MalK | 0.90864 |
| P0A9X9 | Cold shock protein CspA | 0.88728 |
| P0AGG8 | Metalloprotease TldD | 0.88296 |
| P21507 | ATP-dependent RNA helicase SrmB | 0.86637 |
| P0AD61 | Pyruvate kinase I | 0.82680 |
| P14175 | Glycine betaine/proline betaine transport system ATP-binding protein ProV | 0.81530 |
| P0A9D8 | 2,3,4,5-tetrahydropyridine-2,6-dicarboxylate N-succinyltransferase | 0.78766 |
| P00926 | D-serine dehydratase | 0.77116 |
| P37689 | 2,3-bisphosphoglycerate-independent phosphoglycerate mutase | 0.76216 |
| P0A9B2 | Glyceraldehyde-3-phosphate dehydrogenase A | 0.73279 |
| P0A6H5 | ATP-dependent protease ATPase subunit HslU | 0.72846 |
| P0AEU7 | Chaperone protein Skp | 0.72713 |
| P10346 | Glutamine transport ATP-binding protein GlnQ | 0.72150 |
| P0A6Q6 | 3-hydroxyacyl-[acyl-carrier-protein] dehydratase FabZ | 0.71741 |
| P09148 | Galactose-1-phosphate uridylyltransferase | 0.71347 |
| P62707 | 2,3-bisphosphoglycerate-dependent phosphoglycerate mutase | 0.70576 |
| P25536 | dTTP/UTP pyrophosphatase | 0.70364 |
| P0A903 | Outer membrane protein assembly factor BamC | 0.69867 |
| P0A940 | Outer membrane protein assembly factor BamA | 0.69702 |
| P75957 | Lipoprotein-releasing system ATP-binding protein LolD | 0.68717 |
| P0ADE8 | tRNA-modifying protein YgfZ | 0.66981 |
| P0ADS2 | Cell division protein ZapA | 0.66313 |
| P11875 | Arginine--tRNA ligase | 0.65255 |
| P0ACG1 | DNA-binding protein StpA | 0.65248 |
| P0ADA3 | Murein hydrolase activator NlpD | 0.64364 |
| P0AEK2 | 3-oxoacyl-[acyl-carrier-protein] reductase FabG | 0.64253 |
| P0ABQ4 | Dihydrofolate reductase | 0.63197 |
| P0A870 | Transaldolase B | 0.60264 |
| P08622 | Chaperone protein DnaJ | 0.59720 |
| P0ACJ8 | DNA-binding transcriptional dual regulator CRP | 0.59381 |
| P18843 | NH(3)-dependent NAD(+) synthetase | 0.59051 |
| P0ABU2 | Ribosome-binding ATPase YchF | 0.58687 |
| P36680 | Cell division protein ZapD | 0.57671 |
| P0A6H1 | ATP-dependent Clp protease ATP-binding subunit ClpX | 0.57533 |
| P76116 | Uncharacterized protein YncE | 0.56656 |
| P69503 | Adenine phosphoribosyltransferase | 0.55501 |
| P75823 | Low specificity L-threonine aldolase | 0.53672 |
| P0ACC3 | Iron-sulfur cluster insertion protein ErpA | 0.53327 |
| P0AGK4 | RNA-binding protein YhbY | 0.51813 |
| P0ABU0 | 1,4-dihydroxy-2-naphthoyl-CoA synthase | 0.50931 |
| P46853 | Uncharacterized oxidoreductase YhhX | 0.49961 |
| P0AGJ5 | Uncharacterized tRNA/rRNA methyltransferase YfiF | 0.49065 |
| P0A6L0 | Deoxyribose-phosphate aldolase | 0.48947 |
| P0ACA3 | Stringent starvation protein A | 0.47739 |
| P0A9Q5 | Acetyl-coenzyme A carboxylase carboxyl transferase subunit beta | 0.45222 |
| P39406 | Ribosomal RNA small subunit methyltransferase C | 0.44952 |
| P0ACD4 | Iron-sulfur cluster assembly scaffold protein IscU | 0.44507 |
| P0AFY8 | Negative modulator of initiation of replication | 0.44427 |
| P09372 | Protein GrpE | 0.39534 |
| P32695 | tRNA-dihydrouridine(20/20a) synthase | 0.39131 |
| P45577 | RNA chaperone ProQ | 0.38624 |
| P0A744 | Peptide methionine sulfoxide reductase MsrA | 0.36951 |
| P00490 | Maltodextrin phosphorylase | 0.32191 |
| P0A7L3 | 50S ribosomal protein L20 | 0.30289 |
| P0AC41 | Succinate dehydrogenase flavoprotein subunit | 0.24073 |

**Table 5.** **Downregulated proteins at mid-exponential phase (t=3.5 h) in the Δ*sucA* mutant strain relative to the wild type.** FC: Fold Change. A significance threshold of P < 0.05 is applied.

| **Protein ID** | **Description** | **Log_2_(FC)** |
| --- | --- | --- |
| P00968 | Carbamoyl-phosphate synthase large chain | -5.49059 |
| P00805 | L-asparaginase 2 | -5.36069 |
| P39180 | Antigen 43 | -4.83881 |
| P0A7F3 | Aspartate carbamoyltransferase regulatory chain | -4.68139 |
| P0A6F1 | Carbamoyl-phosphate synthase small chain | -4.52087 |
| P0A786 | Aspartate carbamoyltransferase catalytic subunit | -4.15426 |
| P77252 | Uncharacterized protein YkgE | -3.45509 |
| P64463 | Putative selenoprotein YdfZ | -3.11141 |
| P0AFG6 | Dihydrolipoyllysine-residue succinyltransferase component of 2-oxoglutarate dehydrogenase complex | -2.90161 |
| P0A7E1 | Dihydroorotate dehydrogenase (quinone) | -2.83029 |
| P09158 | Polyamine aminopropyltransferase | -2.55574 |
| P31554 | LPS-assembly protein LptD | -2.28723 |
| P77433 | Uncharacterized protein YkgG | -2.13726 |
| P11349 | Respiratory nitrate reductase 1 beta chain | -2.13046 |
| P0ACY9 | Uncharacterized protein YebG | -2.07003 |
| P0A8F0 | Uracil phosphoribosyltransferase | -2.05128 |
| P0ACL9 | Pyruvate dehydrogenase complex repressor | -2.04306 |
| P0A7F6 | S-adenosylmethionine decarboxylase proenzyme | -2.02903 |
| P37646 | Cyclic di-GMP phosphodiesterase PdeH | -2.02673 |
| P24188 | tRNA uridine(34) hydroxylase | -1.95589 |
| P68066 | Autonomous glycyl radical cofactor | -1.92607 |
| P0A998 | Bacterial non-heme ferritin | -1.75740 |
| P64636 | GMP/IMP nucleotidase YrfG | -1.74071 |
| P77609 | Protein FlxA | -1.61383 |
| P0A8R4 | Protein SlyX | -1.61352 |
| P0CG19 | Truncated inactive ribonuclease PH | -1.47597 |
| P69922 | L-fucose isomerase | -1.45884 |
| P77285 | Ion-translocating oxidoreductase complex subunit G | -1.43921 |
| P08244 | Orotidine 5'-phosphate decarboxylase | -1.41115 |
| P0AE48 | Gamma-glutamylcyclotransferase family protein YtfP | -1.38097 |
| P0A6U3 | tRNA uridine 5-carboxymethylaminomethyl modification enzyme MnmG | -1.35702 |
| P36672 | PTS system trehalose-specific EIIBC component | -1.31923 |
| P60340 | tRNA pseudouridine synthase B | -1.28741 |
| P0ABA0 | ATP synthase subunit b | -1.28446 |
| P15639 | Bifunctional purine biosynthesis protein PurH | -1.25320 |
| P75952 | HTH-type transcriptional repressor ComR | -1.23380 |
| P62617 | 2-C-methyl-D-erythritol 2,4-cyclodiphosphate synthase | -1.23321 |
| P33014 | Putative sulfur carrier protein YeeD | -1.20318 |
| P60560 | GMP reductase | -1.18573 |
| P75933 | Flagella basal body P-ring formation protein FlgA | -1.17610 |
| P0AAN3 | Hydrogenase maturation factor HypB | -1.14761 |
| P05020 | Dihydroorotase | -1.12125 |
| P0A7E9 | Uridylate kinase | -1.09479 |
| P0A717 | Ribose-phosphate pyrophosphokinase | -1.09192 |
| P0AF28 | Nitrate/nitrite response regulator protein NarL | -1.04872 |
| P08373 | UDP-N-acetylenolpyruvoylglucosamine reductase | -1.03397 |
| P0A6W9 | Glutamate--cysteine ligase | -1.02245 |
| P0A9P6 | ATP-dependent RNA helicase DeaD | -0.99087 |
| P31802 | Nitrate/nitrite response regulator protein NarP | -0.96777 |
| P0A951 | Spermidine N(1)-acetyltransferase | -0.96401 |
| P0A9C5 | Glutamine synthetase | -0.96040 |
| P25437 | S-(hydroxymethyl)glutathione dehydrogenase | -0.94831 |
| P0A996 | Anaerobic glycerol-3-phosphate dehydrogenase subunit C | -0.89915 |
| P30958 | Transcription-repair-coupling factor | -0.89060 |
| P0AFM6 | Phage shock protein A | -0.88496 |
| P76055 | tRNA-cytidine(32) 2-sulfurtransferase | -0.88050 |
| P32160 | Uncharacterized protein YiiQ | -0.86725 |
| P0A9E5 | Fumarate and nitrate reduction regulatory protein | -0.85817 |
| P0AAD6 | Serine transporter SdaC | -0.85122 |
| P28904 | Trehalose-6-phosphate hydrolase | -0.84059 |
| P0ADG7 | Inosine-5'-monophosphate dehydrogenase | -0.83462 |
| P64534 | Nickel/cobalt homeostasis protein RcnB | -0.83428 |
| P0A993 | Fructose-1,6-bisphosphatase class 1 | -0.83363 |
| P30744 | L-serine dehydratase 2 | -0.82679 |
| P0A9W9 | Protein YrdA | -0.80669 |
| P0ABS8 | DNA polymerase III subunit theta | -0.80347 |
| P0AAT6 | Ribosomal silencing factor RsfS | -0.79274 |
| P0A6I0 | Cytidylate kinase | -0.77758 |
| P56580 | PTS system glucitol/sorbitol-specific EIIB component | -0.77208 |
| P0A9D2 | Glutathione S-transferase GstA | -0.75491 |
| P52627 | Regulator of sigma S factor FliZ | -0.75210 |
| P15640 | Phosphoribosylamine--glycine ligase | -0.75053 |
| P37095 | Peptidase B | -0.73652 |
| P0A707 | Translation initiation factor IF-3 | -0.73217 |
| A0A385XJE6 | Transposase InsH for insertion sequence element IS5U | -0.72291 |
| P14175 | Glycine betaine/proline betaine transport system ATP-binding protein ProV | -0.71933 |
| P0A7I7 | GTP cyclohydrolase-2 | -0.71866 |
| P69222 | Translation initiation factor IF-1 | -0.71672 |
| P0AG82 | Phosphate-binding protein PstS | -0.70514 |
| P69441 | Adenylate kinase | -0.69524 |
| P0AD33 | UPF0381 protein YfcZ | -0.69515 |
| P0A7S9 | 30S ribosomal protein S13 | -0.68355 |
| P23909 | DNA mismatch repair protein MutS | -0.68049 |
| Q46856 | Alcohol dehydrogenase YqhD | -0.66647 |
| P0A7E5 | CTP synthase | -0.66441 |
| P27829 | UDP-N-acetyl-D-mannosamine dehydrogenase | -0.66273 |
| P0ADG4 | Nus factor SuhB | -0.65779 |
| P0A8F4 | Uridine kinase | -0.65773 |
| P06968 | Deoxyuridine 5'-triphosphate nucleotidohydrolase | -0.65508 |
| P31475 | Uncharacterized HTH-type transcriptional regulator YieP | -0.64722 |
| P75838 | Ribosomal protein S12 methylthiotransferase accessory factor YcaO | -0.63099 |
| P0A9P4 | Thioredoxin reductase | -0.61976 |
| P60716 | Lipoyl synthase | -0.60926 |
| P0A8H6 | Der GTPase-activating protein YihI | -0.60123 |
| P0A6N4 | Elongation factor P | -0.59263 |
| P0A7I4 | Peptide chain release factor RF3 | -0.59056 |
| P0ABR1 | DNA damage-inducible protein I | -0.58973 |
| P0AEI4 | Ribosomal protein S12 methylthiotransferase RimO | -0.58184 |
| P17117 | Oxygen-insensitive NADPH nitroreductase | -0.57469 |
| P21888 | Cysteine--tRNA ligase | -0.57240 |
| P76658 | Bifunctional protein HldE | -0.56685 |
| P0A7D7 | Phosphoribosylaminoimidazole-succinocarboxamide synthase | -0.55391 |
| P0A840 | 5'/3'-nucleotidase SurE | -0.54029 |
| P0ACR9 | Transcriptional repressor MprA | -0.50187 |
| P0A7N9 | 50S ribosomal protein L33 | -0.49347 |
| P32695 | tRNA-dihydrouridine(20/20a) synthase | -0.48001 |
| P0A6N8 | Elongation factor P-like protein | -0.47260 |
| P09030 | Exodeoxyribonuclease III | -0.46453 |
| P65556 | Uncharacterized Nudix hydrolase YfcD | -0.46183 |
| P0A7L3 | 50S ribosomal protein L20 | -0.45455 |
| P0AAI5 | 3-oxoacyl-[acyl-carrier-protein] synthase 2 | -0.45346 |
| P0A9K9 | FKBP-type peptidyl-prolyl cis-trans isomerase SlyD | -0.45337 |
| P23893 | Glutamate-1-semialdehyde 2,1-aminomutase | -0.44380 |
| P0A9L5 | Peptidyl-prolyl cis-trans isomerase C | -0.44022 |
| P11875 | Arginine--tRNA ligase | -0.43247 |
| P37744 | Glucose-1-phosphate thymidylyltransferase 1 | -0.43235 |
| P39199 | 50S ribosomal protein L3 glutamine methyltransferase | -0.42720 |
| P21599 | Pyruvate kinase II | -0.42565 |
| P0A6R0 | Beta-ketoacyl-[acyl-carrier-protein] synthase III | -0.40053 |
| P0A749 | UDP-N-acetylglucosamine 1-carboxyvinyltransferase | -0.39039 |
| P60624 | 50S ribosomal protein L24 | -0.38563 |
| P0ABS1 | RNA polymerase-binding transcription factor DksA | -0.38404 |
| P0A800 | DNA-directed RNA polymerase subunit omega | -0.37378 |
| P0A6A3 | Acetate kinase | -0.37266 |
| P0A850 | Trigger factor | -0.35233 |
| P07017 | Methyl-accepting chemotaxis protein II | -0.34303 |
| P42641 | GTPase ObgE/CgtA | -0.32845 |
| P0ABQ4 | Dihydrofolate reductase | -0.31487 |
| P0A9X4 | Cell shape-determining protein MreB | -0.30732 |
| P0A6P1 | Elongation factor Ts | -0.30448 |
| P0A9C9 | Fructose-1,6-bisphosphatase 1 class 2 | -0.29873 |
| P0AA10 | 50S ribosomal protein L13 | -0.29648 |
| P0A817 | S-adenosylmethionine synthase | -0.29281 |
| P0A6D7 | Shikimate kinase 1 | -0.28170 |
| P37745 | dTDP-4-dehydrorhamnose 3,5-epimerase | -0.28083 |
| P0A7W7 | 30S ribosomal protein S8 | -0.27168 |
| P0A8A2 | Probable transcriptional regulatory protein YeeN | -0.26264 |
| P60651 | Agmatinase | -0.24632 |
| P02359 | 30S ribosomal protein S7 | -0.23205 |
| P0A7W1 | 30S ribosomal protein S5 | -0.22901 |
| P0AFG0 | Transcription termination/antitermination protein NusG | -0.21786 |

**Table 6. Downregulated proteins at mid-exponential phase (t=3.5 h) in the Δ*gltA* mutant strain relative to the wild type.** FC: Fold Change. A significance threshold of P < 0.05 is applied.

| **Protein ID** | **Description** | **Log_2_(FC)** |
| --- | --- | --- |
| P39180 | Antigen 43 | -3.45274 |
| P30958 | Transcription-repair-coupling factor | -2.15142 |
| P64463 | Putative selenoprotein YdfZ | -1.95874 |
| P0ACP7 | HTH-type transcriptional repressor PurR | -1.78709 |
| P09158 | Polyamine aminopropyltransferase | -1.78241 |
| P00805 | L-asparaginase 2 | -1.71077 |
| P00393 | Type II NADH:quinone oxidoreductase | -1.68217 |
| P37750 | Putative lipopolysaccharide biosynthesis O-acetyl transferase WbbJ | -1.61251 |
| P69922 | L-fucose isomerase | -1.41913 |
| P0A7S3 | 30S ribosomal protein S12 | -1.41352 |
| P0A9F3 | HTH-type transcriptional regulator CysB | -1.41027 |
| P08373 | UDP-N-acetylenolpyruvoylglucosamine reductase | -1.33756 |
| P0A6U5 | Ribosomal RNA small subunit methyltransferase G | -1.28667 |
| P0ADY7 | 50S ribosomal protein L16 | -1.27439 |
| P11349 | Respiratory nitrate reductase 1 beta chain | -1.23894 |
| P0A6I6 | Phosphopantetheine adenylyltransferase | -1.15003 |
| P24188 | tRNA uridine(34) hydroxylase | -1.12152 |
| P52627 | Regulator of sigma S factor FliZ | -1.06550 |
| P77756 | 7-cyano-7-deazaguanine synthase | -1.02836 |
| P0A7L8 | 50S ribosomal protein L27 | -1.00686 |
| P68066 | Autonomous glycyl radical cofactor | -0.98759 |
| P0ABH0 | Cell division protein FtsA | -0.98156 |
| P00363 | Fumarate reductase flavoprotein subunit | -0.98031 |
| P0ACR9 | Transcriptional repressor MprA | -0.95938 |
| P0A884 | Thymidylate synthase | -0.95598 |
| P0A993 | Fructose-1,6-bisphosphatase class 1 | -0.95384 |
| P0AGF2 | Sulfur acceptor protein CsdE | -0.94553 |
| P77609 | Protein FlxA | -0.94087 |
| P75933 | Flagella basal body P-ring formation protein FlgA | -0.93881 |
| Q46856 | Alcohol dehydrogenase YqhD | -0.93463 |
| P09373 | Formate acetyltransferase 1 | -0.91198 |
| P0AES4 | DNA gyrase subunit A | -0.89360 |
| P0A7X3 | 30S ribosomal protein S9 | -0.87921 |
| P0A998 | Bacterial non-heme ferritin | -0.86249 |
| P21889 | Aspartate--tRNA ligase | -0.84916 |
| P0AE78 | Magnesium and cobalt efflux protein CorC | -0.83785 |
| P05707 | Sorbitol-6-phosphate 2-dehydrogenase | -0.81015 |
| P0A7V8 | 30S ribosomal protein S4 | -0.80757 |
| P37349 | PEP-dependent dihydroxyacetone kinase, phosphoryl donor subunit DhaM | -0.80706 |
| P0AD07 | Uncharacterized protein YecF | -0.80578 |
| P69874 | Spermidine/putrescine import ATP-binding protein PotA | -0.80242 |
| P0AAT6 | Ribosomal silencing factor RsfS | -0.77384 |
| P0A7E1 | Dihydroorotate dehydrogenase (quinone) | -0.75333 |
| P00934 | Threonine synthase | -0.71580 |
| P37759 | dTDP-glucose 4,6-dehydratase 1 | -0.69595 |
| P76015 | PEP-dependent dihydroxyacetone kinase, dihydroxyacetone-binding subunit DhaK | -0.69354 |
| P0A8F0 | Uracil phosphoribosyltransferase | -0.67256 |
| P00968 | Carbamoyl-phosphate synthase large chain | -0.66807 |
| P05852 | tRNA N6-adenosine threonylcarbamoyltransferase | -0.66564 |
| P64636 | GMP/IMP nucleotidase YrfG | -0.65873 |
| P0AEI4 | Ribosomal protein S12 methylthiotransferase RimO | -0.65401 |
| P75960 | NAD-dependent protein deacylase | -0.65031 |
| P0A9N4 | Pyruvate formate-lyase 1-activating enzyme | -0.64945 |
| P33920 | Nucleoid-associated protein YejK | -0.64770 |
| P0A7V3 | 30S ribosomal protein S3 | -0.64607 |
| P0A7S9 | 30S ribosomal protein S13 | -0.64103 |
| P0A7I4 | Peptide chain release factor RF3 | -0.63082 |
| P0A7U7 | 30S ribosomal protein S20 | -0.62794 |
| P0A7K6 | 50S ribosomal protein L19 | -0.61754 |
| P21888 | Cysteine--tRNA ligase | -0.60973 |
| P0A8T1 | Ribosomal protein L11 methyltransferase | -0.59937 |
| P0AC53 | Glucose-6-phosphate 1-dehydrogenase | -0.59082 |
| P00961 | Glycine--tRNA ligase beta subunit | -0.57546 |
| P0A7R9 | 30S ribosomal protein S11 | -0.57499 |
| P0A951 | Spermidine N(1)-acetyltransferase | -0.56328 |
| P0A707 | Translation initiation factor IF-3 | -0.54739 |
| P0A7E5 | CTP synthase | -0.54241 |
| P05706 | PTS system glucitol/sorbitol-specific EIIA component | -0.51229 |
| P0CG19 | Truncated inactive ribonuclease PH | -0.50219 |
| P62623 | 4-hydroxy-3-methylbut-2-enyl diphosphate reductase | -0.47358 |
| P0A6Z1 | Chaperone protein HscA | -0.46744 |
| P09147 | UDP-glucose 4-epimerase | -0.45891 |
| P0ABF6 | Cytidine deaminase | -0.44702 |
| P0A6R0 | Beta-ketoacyl-[acyl-carrier-protein] synthase III | -0.44660 |
| P36680 | Cell division protein ZapD | -0.43517 |
| P0A6P9 | Enolase | -0.43438 |
| P42641 | GTPase ObgE/CgtA | -0.43176 |
| P0ACG1 | DNA-binding protein StpA | -0.41305 |
| P0A7M2 | 50S ribosomal protein L28 | -0.41175 |
| P31802 | Nitrate/nitrite response regulator protein NarP | -0.40446 |
| P0AC47 | Fumarate reductase iron-sulfur subunit | -0.39820 |
| P37744 | Glucose-1-phosphate thymidylyltransferase 1 | -0.39635 |
| P37745 | dTDP-4-dehydrorhamnose 3,5-epimerase | -0.39617 |
| P23893 | Glutamate-1-semialdehyde 2,1-aminomutase | -0.37193 |
| P08839 | Phosphoenolpyruvate-protein phosphotransferase | -0.36939 |
| P0A749 | UDP-N-acetylglucosamine 1-carboxyvinyltransferase | -0.36772 |
| P26646 | Probable acrylyl-CoA reductase AcuI | -0.36581 |
| P0ACA3 | Stringent starvation protein A | -0.36175 |
| P0ADG7 | Inosine-5'-monophosphate dehydrogenase | -0.35725 |
| P0A7T3 | 30S ribosomal protein S16 | -0.35585 |
| P16659 | Proline--tRNA ligase | -0.35288 |
| P0AF08 | Iron-sulfur cluster carrier protein | -0.35239 |
| P00946 | Mannose-6-phosphate isomerase | -0.34433 |
| P38521 | Uncharacterized protein YggL | -0.33403 |
| P25437 | S-(hydroxymethyl)glutathione dehydrogenase | -0.33228 |
| P52614 | Flagellar hook-length control protein | -0.32566 |
| P68919 | 50S ribosomal protein L25 | -0.32336 |
| P0A7N4 | 50S ribosomal protein L32 | -0.31534 |
| P0A6A3 | Acetate kinase | -0.31426 |
| P15288 | Cytosol non-specific dipeptidase | -0.31385 |
| P0AB18 | Sulfurtransferase TusE | -0.31128 |
| P37689 | 2,3-bisphosphoglycerate-independent phosphoglycerate mutase | -0.30751 |
| P0A817 | S-adenosylmethionine synthase | -0.27384 |
| P02359 | 30S ribosomal protein S7 | -0.26915 |
| P30850 | Exoribonuclease 2 | -0.26172 |
| P0AEP3 | UTP--glucose-1-phosphate uridylyltransferase | -0.25936 |
| P39173 | Putative glucose-6-phosphate 1-epimerase | -0.25420 |
| P0A9X4 | Cell shape-determining protein MreB | -0.24925 |
| P0A717 | Ribose-phosphate pyrophosphokinase | -0.24827 |
| P0A6P1 | Elongation factor Ts | -0.22376 |
| P0A7X6 | Ribosome maturation factor RimM | -0.20019 |
| P0A870 | Transaldolase B | -0.19840 |
| P0A6D7 | Shikimate kinase 1 | -0.19126 |
| P0A8F4 | Uridine kinase | -0.18706 |
| P0ABS1 | RNA polymerase-binding transcription factor DksA | -0.17081 |
| P0AF12 | 5'-methylthioadenosine/S-adenosylhomocysteine nucleosidase | -0.12218 |

**Table 7. Downregulated proteins at mid-exponential phase (t=3.5 h) in the Δ*nuoI* mutant strain relative to the wild type.** FC: Fold Change. A significance threshold of P < 0.05 is applied.

| **Protein ID** | **Description** | **Log_2_(FC)** |
| --- | --- | --- |
| P39180 | Antigen 43 | -3.28283 |
| P09158 | Polyamine aminopropyltransferase | -3.13307 |
| P64463 | Putative selenoprotein YdfZ | -2.53986 |
| P00582 | DNA polymerase I | -2.41479 |
| P31554 | LPS-assembly protein LptD | -2.37682 |
| P37646 | Cyclic di-GMP phosphodiesterase PdeH | -2.23275 |
| P00968 | Carbamoyl-phosphate synthase large chain | -2.09876 |
| P08244 | Orotidine 5'-phosphate decarboxylase | -2.07320 |
| P0A847 | Queuine tRNA-ribosyltransferase | -2.03487 |
| P0A7M6 | 50S ribosomal protein L29 | -1.98773 |
| P22106 | Asparagine synthetase B [glutamine-hydrolyzing] | -1.92762 |
| P63389 | Probable ATP-binding protein YheS | -1.85126 |
| P08373 | UDP-N-acetylenolpyruvoylglucosamine reductase | -1.83857 |
| P00393 | Type II NADH:quinone oxidoreductase | -1.75058 |
| P33920 | Nucleoid-associated protein YejK | -1.73626 |
| P0AFM2 | Glycine betaine/proline betaine-binding periplasmic protein | -1.68732 |
| P21170 | Biosynthetic arginine decarboxylase | -1.55339 |
| P0A8F0 | Uracil phosphoribosyltransferase | -1.52927 |
| P0A993 | Fructose-1,6-bisphosphatase class 1 | -1.52288 |
| P0A6I6 | Phosphopantetheine adenylyltransferase | -1.51819 |
| P0A7E1 | Dihydroorotate dehydrogenase (quinone) | -1.48847 |
| P0A6T5 | GTP cyclohydrolase 1 | -1.47257 |
| P23538 | Phosphoenolpyruvate synthase | -1.41285 |
| P0A7S3 | 30S ribosomal protein S12 | -1.40075 |
| P0ACQ4 | Hydrogen peroxide-inducible genes activator | -1.38000 |
| P0A6F1 | Carbamoyl-phosphate synthase small chain | -1.36595 |
| P0AG59 | 30S ribosomal protein S14 | -1.35235 |
| P64455 | Uncharacterized protein YdcY | -1.34858 |
| P0C058 | Small heat shock protein IbpB | -1.24523 |
| P0A9C5 | Glutamine synthetase | -1.21171 |
| P0A7F3 | Aspartate carbamoyltransferase regulatory chain | -1.20419 |
| P0A6U5 | Ribosomal RNA small subunit methyltransferase G | -1.20310 |
| P23869 | Peptidyl-prolyl cis-trans isomerase B | -1.15420 |
| P0A884 | Thymidylate synthase | -1.14321 |
| P37759 | dTDP-glucose 4,6-dehydratase 1 | -1.13569 |
| P64636 | GMP/IMP nucleotidase YrfG | -1.11679 |
| P07363 | Chemotaxis protein CheA | -1.03335 |
| P0AES6 | DNA gyrase subunit B | -1.03093 |
| P0A9N4 | Pyruvate formate-lyase 1-activating enzyme | -0.99788 |
| P0AE78 | Magnesium and cobalt efflux protein CorC | -0.99656 |
| P0AES4 | DNA gyrase subunit A | -0.96236 |
| P00452 | Ribonucleoside-diphosphate reductase 1 subunit alpha | -0.93215 |
| P0AF90 | Regulator of ribonuclease activity B | -0.91591 |
| P31120 | Phosphoglucosamine mutase | -0.89083 |
| P0AEQ3 | Glutamine-binding periplasmic protein | -0.88638 |
| P21889 | Aspartate--tRNA ligase | -0.83880 |
| P0CG19 | Truncated inactive ribonuclease PH | -0.83563 |
| P00363 | Fumarate reductase flavoprotein subunit | -0.82765 |
| Q46857 | 2,5-diketo-D-gluconic acid reductase A | -0.82549 |
| P0A8W0 | HTH-type transcriptional repressor NanR | -0.82359 |
| P60560 | GMP reductase | -0.79829 |
| P05852 | tRNA N6-adenosine threonylcarbamoyltransferase | -0.76982 |
| P0A9F3 | HTH-type transcriptional regulator CysB | -0.75402 |
| P37349 | PEP-dependent dihydroxyacetone kinase, phosphoryl donor subunit DhaM | -0.73657 |
| P00934 | Threonine synthase | -0.72460 |
| P77609 | Protein FlxA | -0.71258 |
| P77552 | Uncharacterized protein YdhQ | -0.63996 |
| P00448 | Superoxide dismutase [Mn] | -0.63837 |
| P0ADX7 | Uncharacterized protein YhhA | -0.63170 |
| P00961 | Glycine--tRNA ligase beta subunit | -0.63006 |
| Q46856 | Alcohol dehydrogenase YqhD | -0.62561 |
| P0A9P6 | ATP-dependent RNA helicase DeaD | -0.60428 |
| P0A8R4 | Protein SlyX | -0.59727 |
| P0A7S9 | 30S ribosomal protein S13 | -0.57939 |
| P62623 | 4-hydroxy-3-methylbut-2-enyl diphosphate reductase | -0.57491 |
| P0A7T3 | 30S ribosomal protein S16 | -0.57371 |
| P0A7K6 | 50S ribosomal protein L19 | -0.56296 |
| P27838 | Iron-sulfur cluster assembly protein CyaY | -0.54627 |
| P0A7L3 | 50S ribosomal protein L20 | -0.53134 |
| P21338 | Ribonuclease I | -0.46482 |
| P0AEM9 | L-cystine-binding protein TcyJ | -0.44439 |
| P0ABS1 | RNA polymerase-binding transcription factor DksA | -0.38741 |
| P0A7N9 | 50S ribosomal protein L33 | -0.37162 |
| P30850 | Exoribonuclease 2 | -0.34849 |
| P68919 | 50S ribosomal protein L25 | -0.22503 |

**Table 8. Downregulated proteins at mid-exponential phase (t=3.5 h) in the Δ*icd* mutant strain relative to the wild type.** FC: Fold Change. A significance threshold of P < 0.05 is applied.

| **Protein ID** | **Description** | **Log_2_(FC)** |
| --- | --- | --- |
| P11349 | Respiratory nitrate reductase 1 beta chain | -4.89854 |
| P39180 | Antigen 43 | -4.77996 |
| P30958 | Transcription-repair-coupling factor | -4.34835 |
| P76010 | Flagellar brake protein YcgR | -4.20928 |
| P00968 | Carbamoyl-phosphate synthase large chain | -3.77696 |
| P0A7F3 | Aspartate carbamoyltransferase regulatory chain | -3.45134 |
| P37646 | Cyclic di-GMP phosphodiesterase PdeH | -3.26367 |
| P0A6F1 | Carbamoyl-phosphate synthase small chain | -3.24331 |
| P31554 | LPS-assembly protein LptD | -3.20273 |
| P09546 | Bifunctional protein PutA | -3.15245 |
| P38038 | Sulfite reductase [NADPH] flavoprotein alpha-component | -3.12854 |
| P68066 | Autonomous glycyl radical cofactor | -3.03245 |
| P76297 | Flagellar protein FlhE | -2.97074 |
| P23538 | Phosphoenolpyruvate synthase | -2.93409 |
| P64463 | Putative selenoprotein YdfZ | -2.89096 |
| P0A786 | Aspartate carbamoyltransferase catalytic subunit | -2.73188 |
| P77285 | Ion-translocating oxidoreductase complex subunit G | -1.97923 |
| P0A7E1 | Dihydroorotate dehydrogenase (quinone) | -1.97408 |
| P07363 | Chemotaxis protein CheA | -1.76012 |
| P0A9H7 | Cyclopropane-fatty-acyl-phospholipid synthase | -1.75837 |
| P09158 | Polyamine aminopropyltransferase | -1.73733 |
| P0C079 | Antitoxin RelB | -1.60867 |
| P0A8F0 | Uracil phosphoribosyltransferase | -1.58904 |
| P0A908 | MltA-interacting protein | -1.45106 |
| P0AAN3 | Hydrogenase maturation factor HypB | -1.12085 |
| P60560 | GMP reductase | -1.03262 |
| P0A8R4 | Protein SlyX | -0.97307 |
| P0AED5 | Response regulator UvrY | -0.94106 |
| P08244 | Orotidine 5'-phosphate decarboxylase | -0.92134 |
| P26616 | NAD-dependent malic enzyme | -0.90084 |
| P37095 | Peptidase B | -0.88478 |
| P09373 | Formate acetyltransferase 1 | -0.83885 |
| P31802 | Nitrate/nitrite response regulator protein NarP | -0.78530 |
| P0ABK5 | Cysteine synthase A | -0.74912 |
| P33136 | Glucans biosynthesis protein G | -0.62861 |
| P33599 | NADH-quinone oxidoreductase subunit C/D | -0.60479 |
| P0ABS8 | DNA polymerase III subunit theta | -0.55327 |
| P0A853 | Tryptophanase | -0.54866 |
| P64455 | Uncharacterized protein YdcY | -0.54733 |
| P62617 | 2-C-methyl-D-erythritol 2,4-cyclodiphosphate synthase | -0.41459 |
| P27838 | Iron-sulfur cluster assembly protein CyaY | -0.32192 |
| P0C0R7 | Ribosomal RNA large subunit methyltransferase E | -0.27202 |
| P0ABD3 | Bacterioferritin | -0.23494 |

**Table 9.** **The pathway analysis for the upregulated proteins in the Δ*sucA* strain compared to the wild type.** This analysis integrates statistical analysis across the entire genome and includes various functional pathway classification frameworks such as Gene Ontology annotations, KEGG pathways, Uniprot, and STRING. ***Count in Network:*** The first number indicates how many proteins in our network are annotated with a particular term. The second number indicates how many proteins in total (in our network and the background) have this term assigned. ***Strength:*** Log10(observed/expected). This measure describes how large the enrichment effect is. It’s the ratio between i) the number of proteins in our network that are annotated with a term and ii) the number of proteins that we expect to be annotated with this term in a random network of the same size. ***False Discovery Rate:*** This measure describes how significant the enrichment is. Shown are P-values corrected for multiple testing within each category using the Benjamini–Hochberg procedure.

Note: When discussing our findings in the manuscript, we primarily reference the local network cluster (STRING), as it offers the advantage of broader coverage, including potential novel modules that might not yet be classified as pathways.

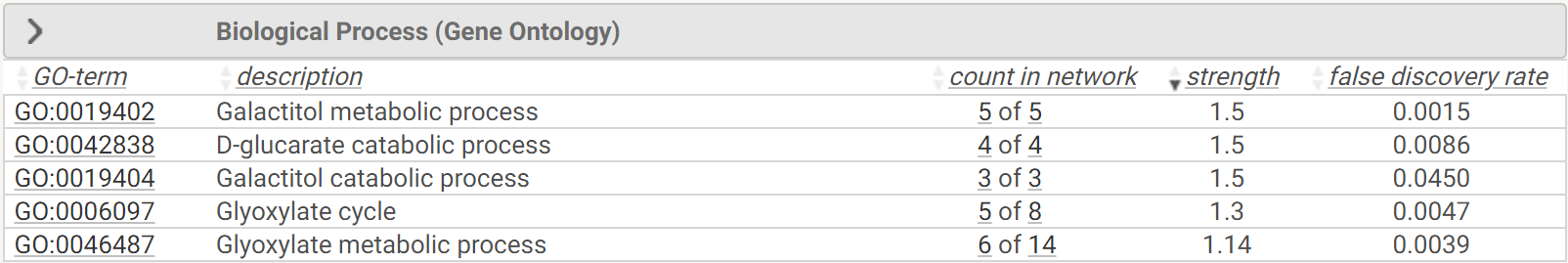

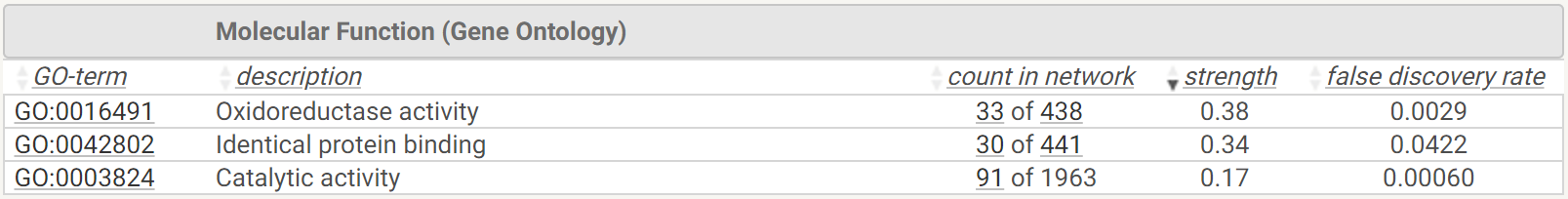

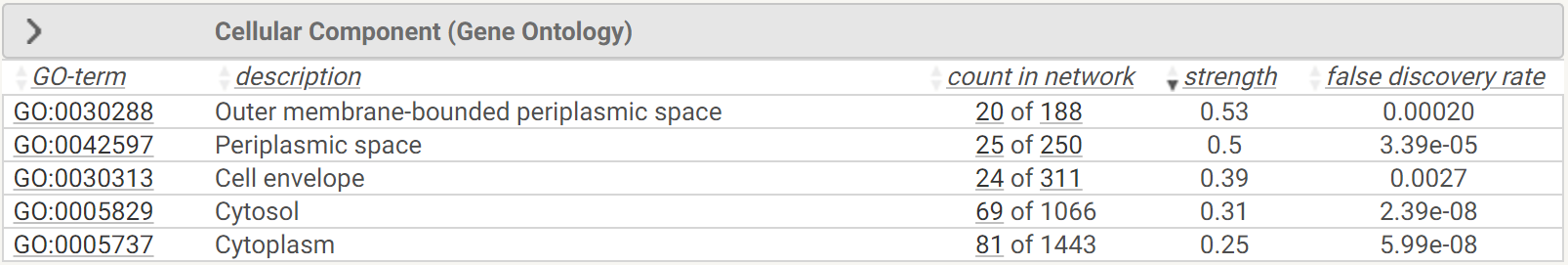

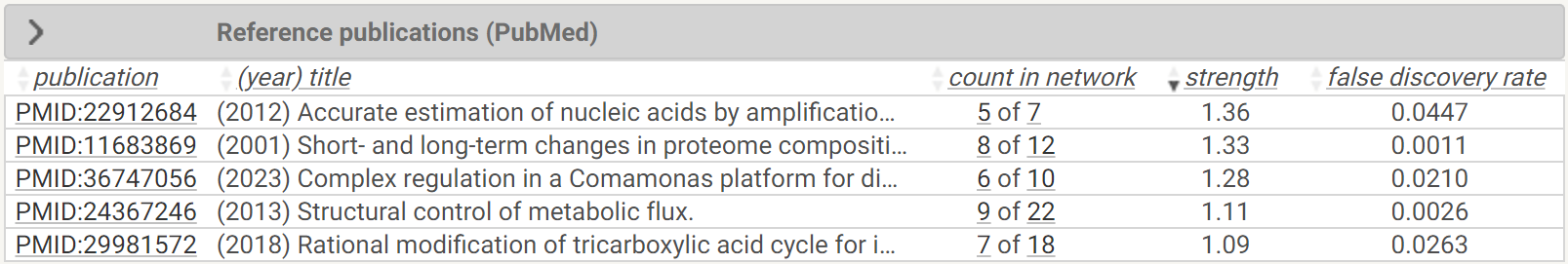

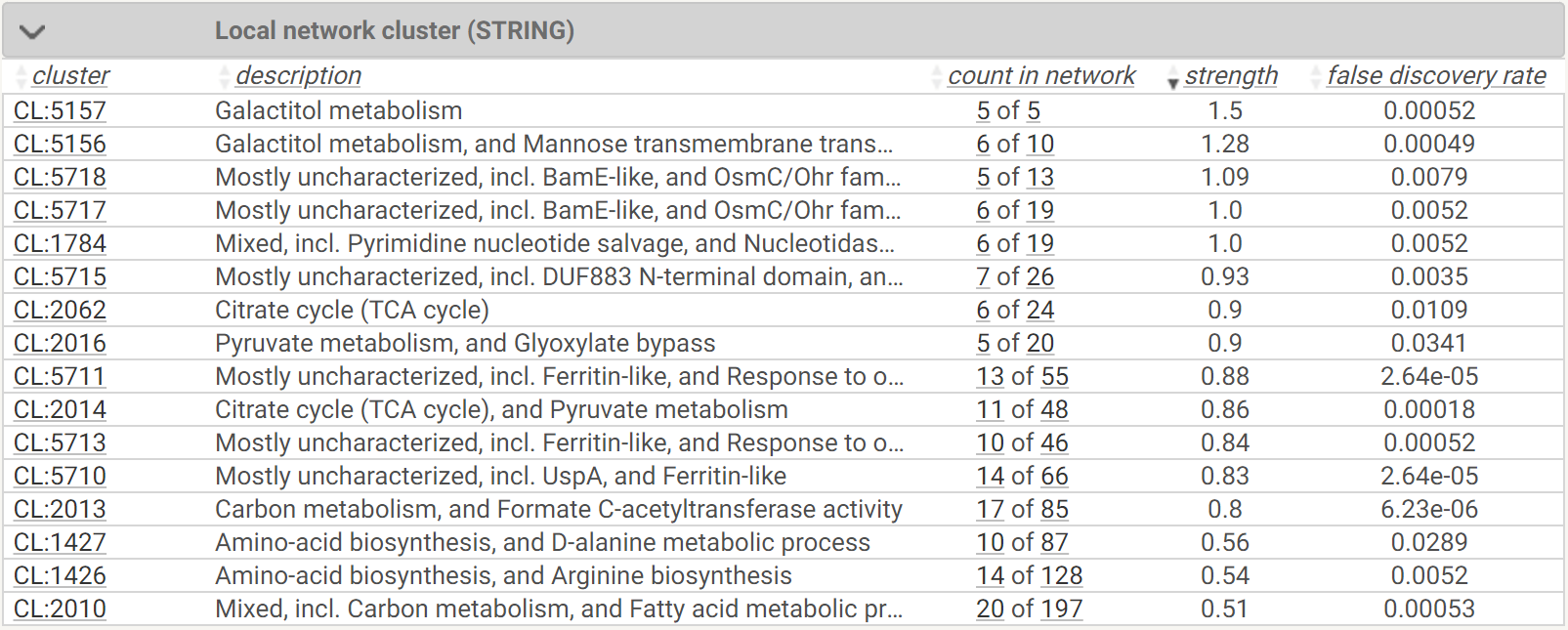

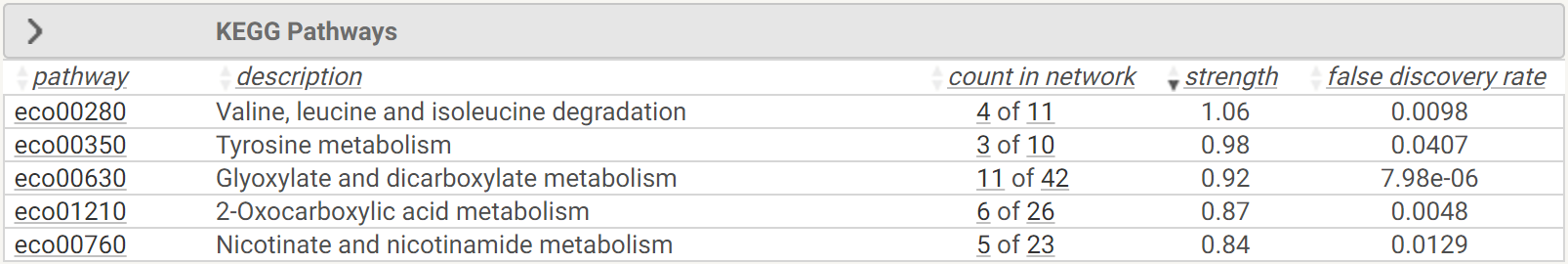

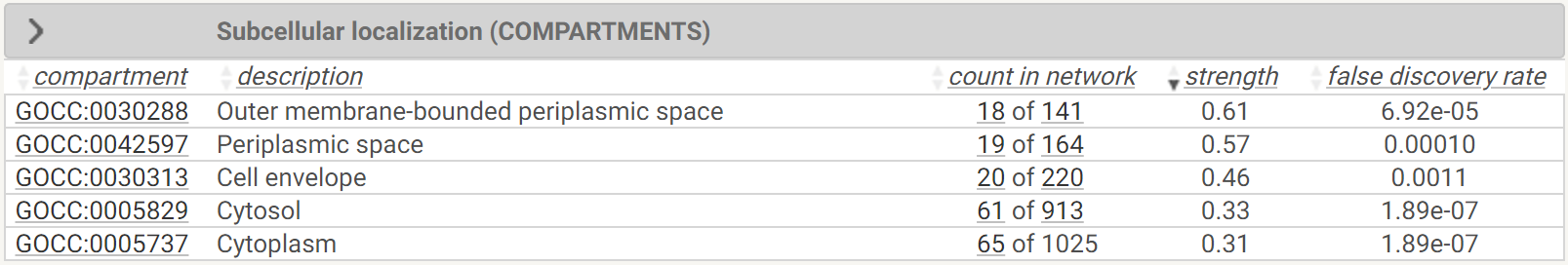

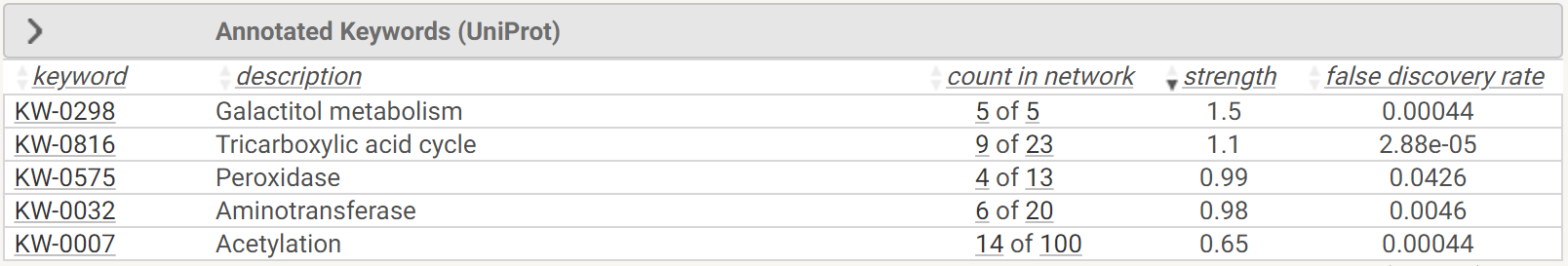

**Table 10.** **The pathway analysis for the upregulated proteins in the Δ*gltA* strain compared to the wild type.** See the legend of Table 9 for further details.

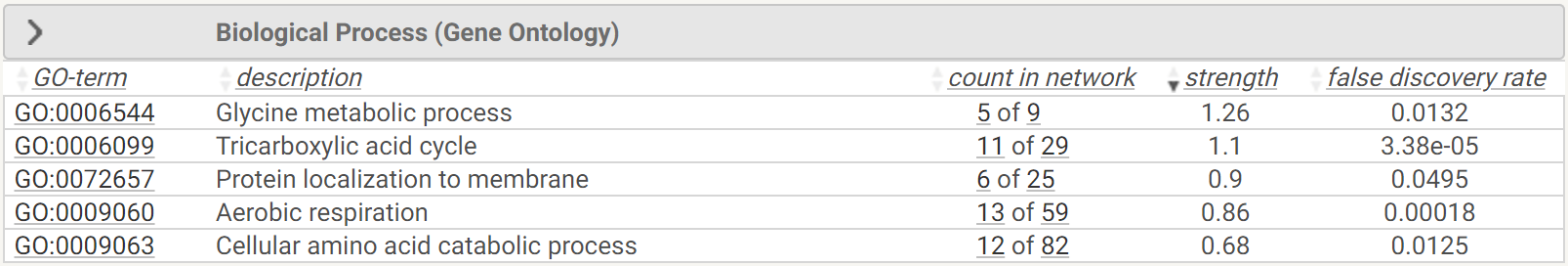

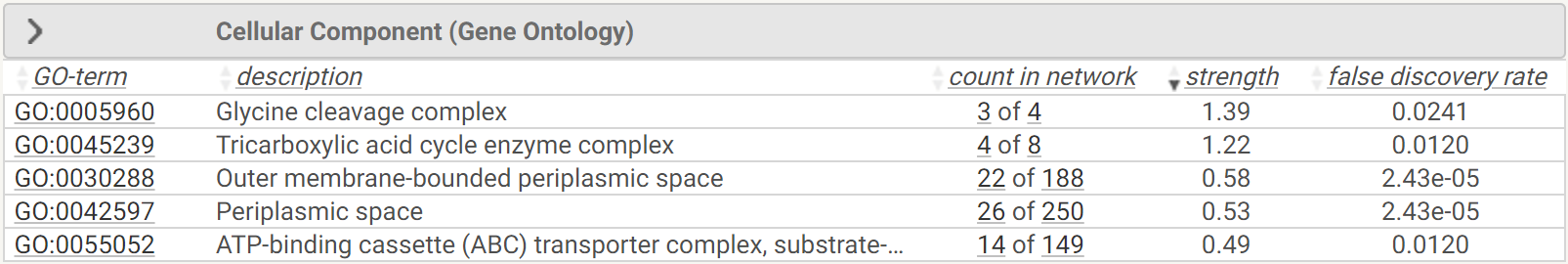

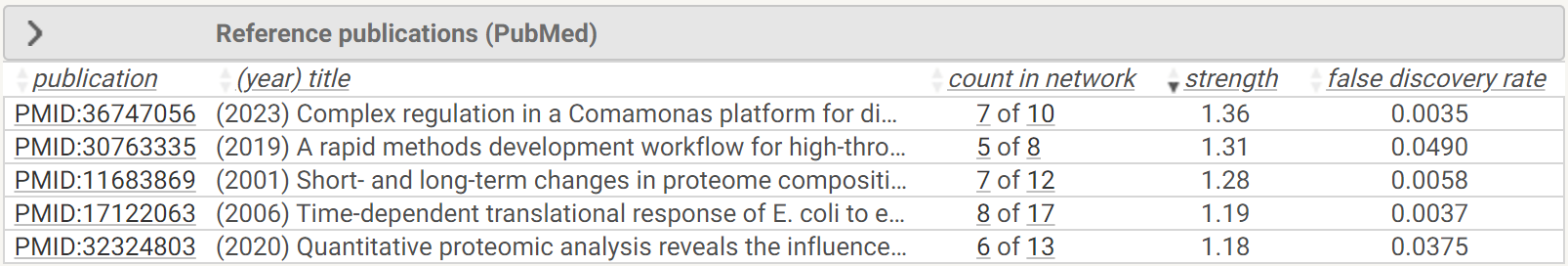

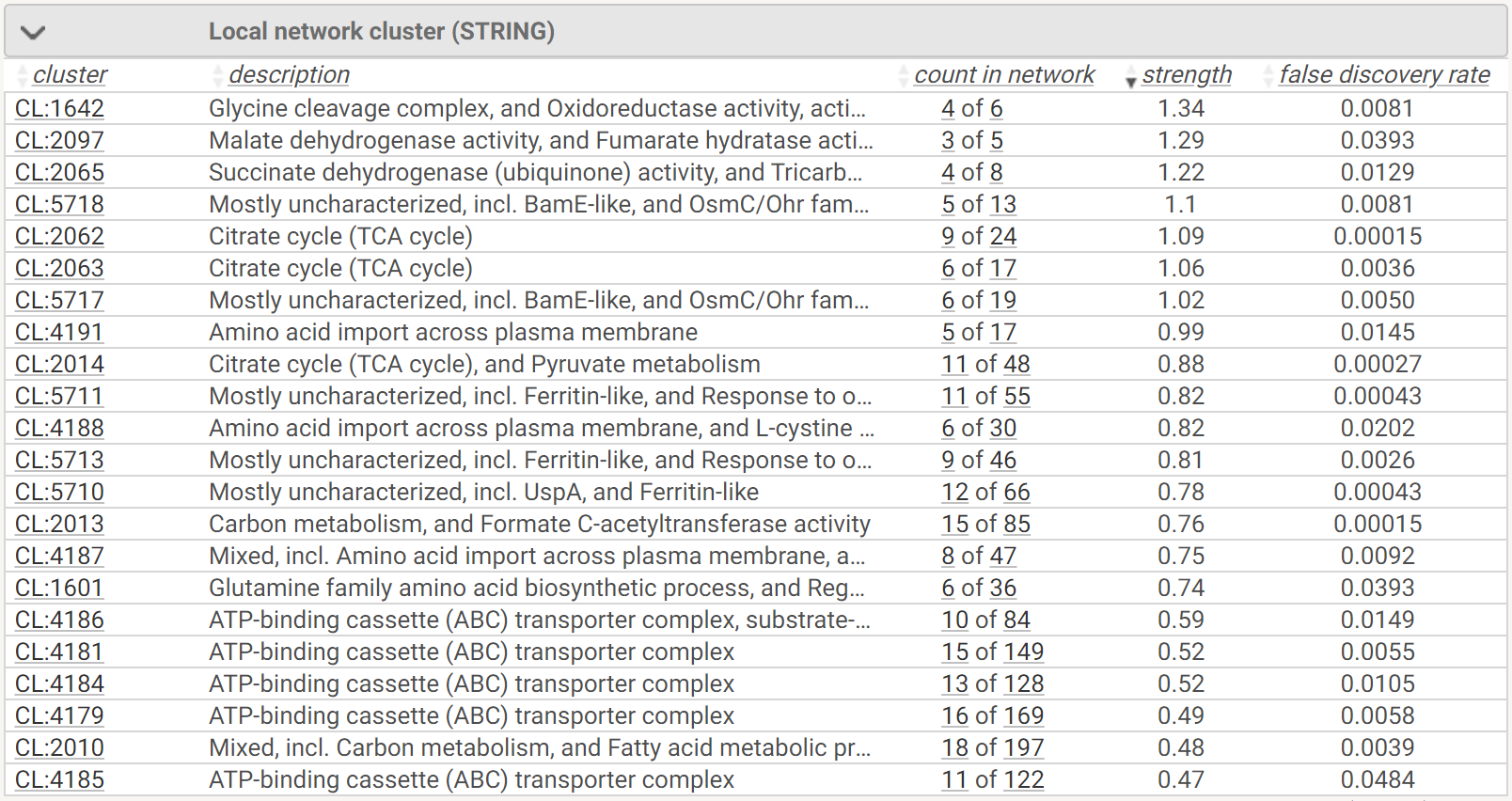

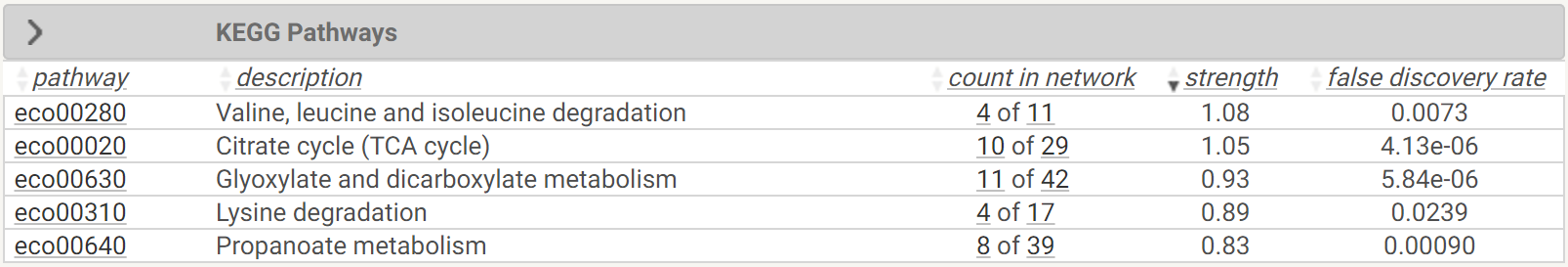

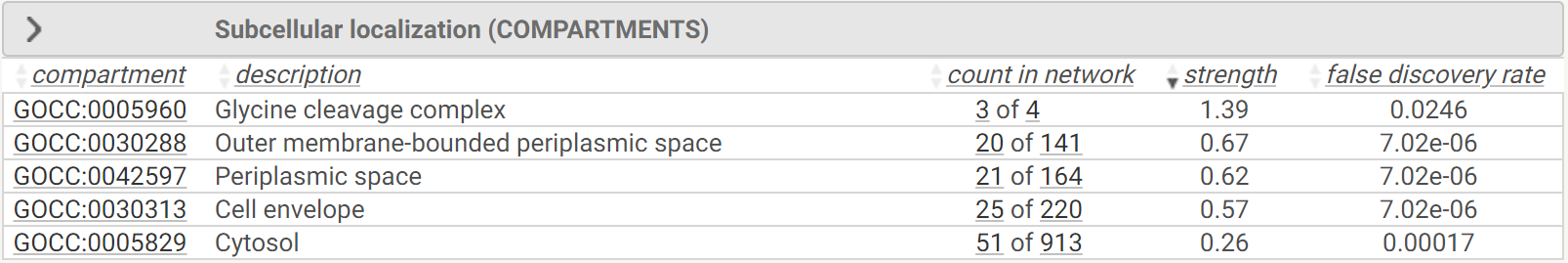

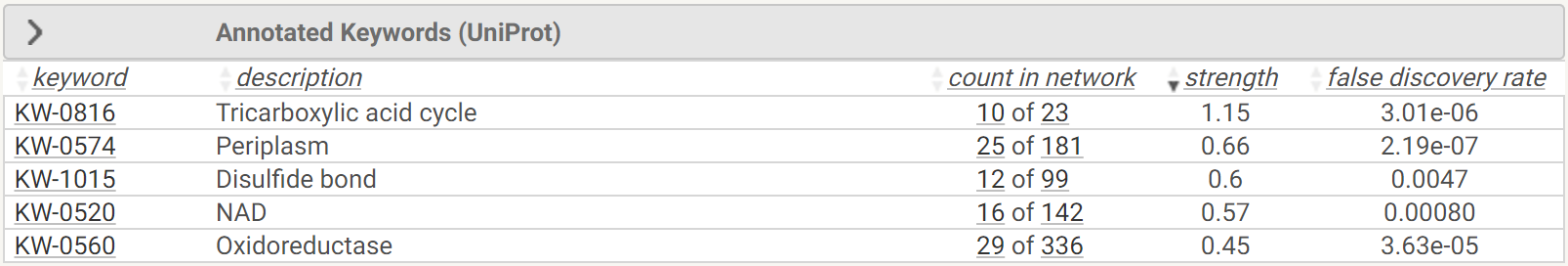

**Table 11.** **The pathway analysis for the upregulated proteins in the Δ*nuoI* strain compared to the wild type.** See the legend of Table 9 for further details.

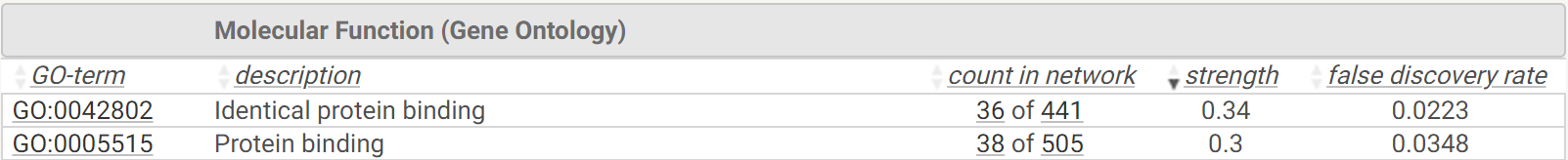

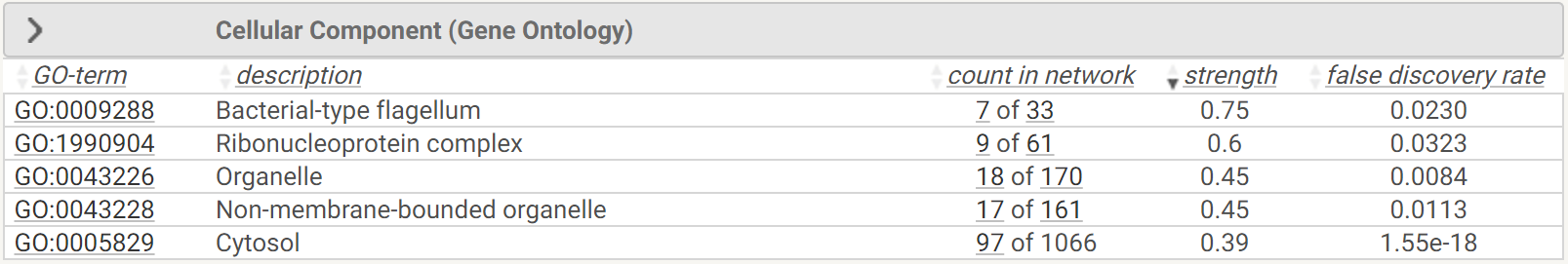

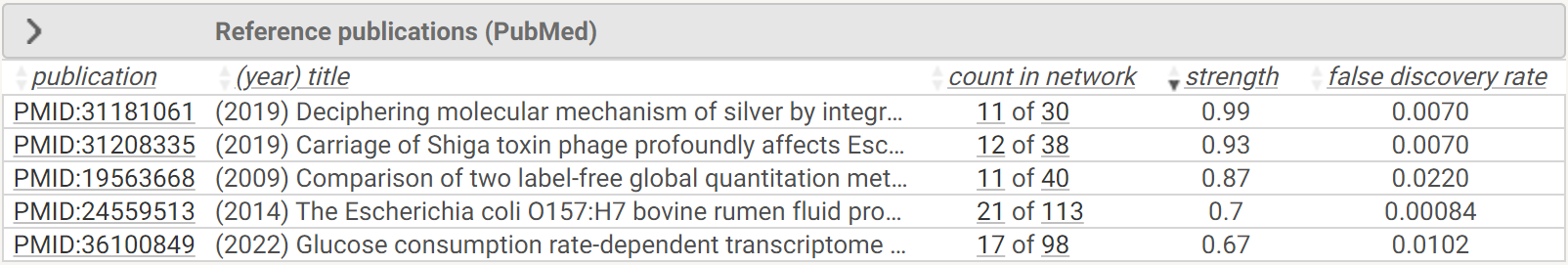

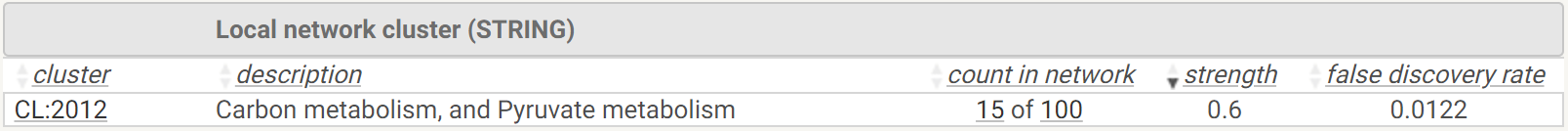

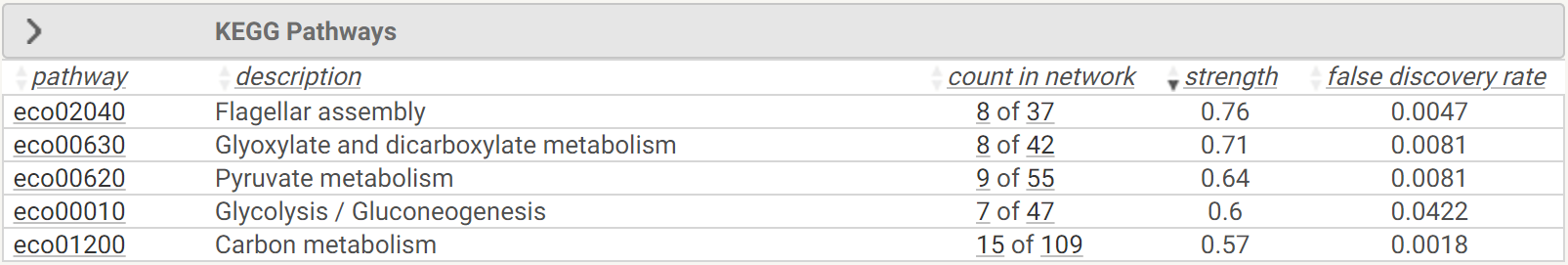

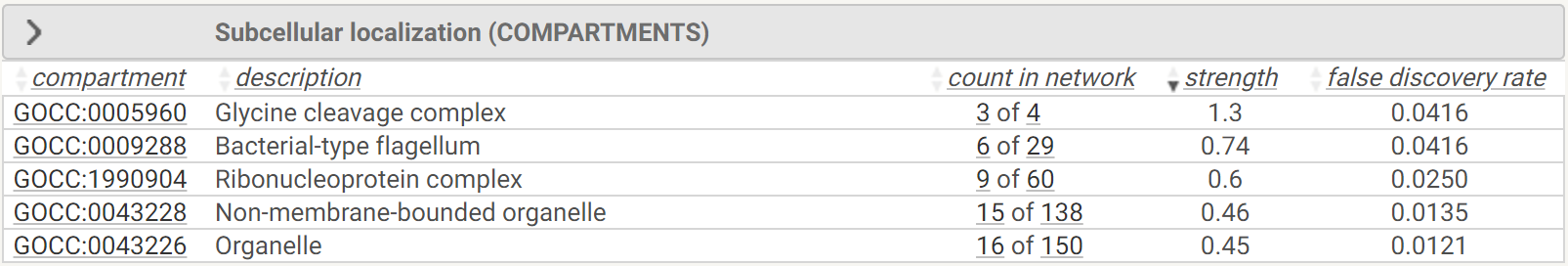

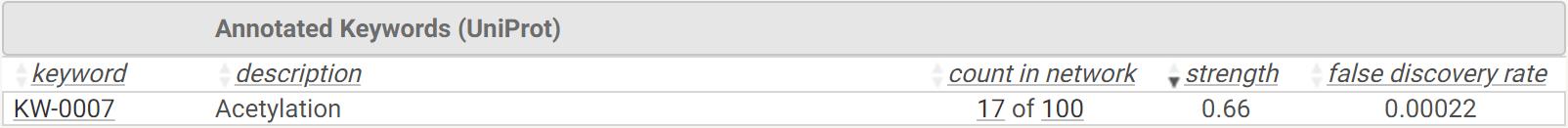

**Table 12.** **The pathway analysis for the upregulated proteins in the Δ*icd* strain compared to the wild type.** See the legend of Table 9 for further details.

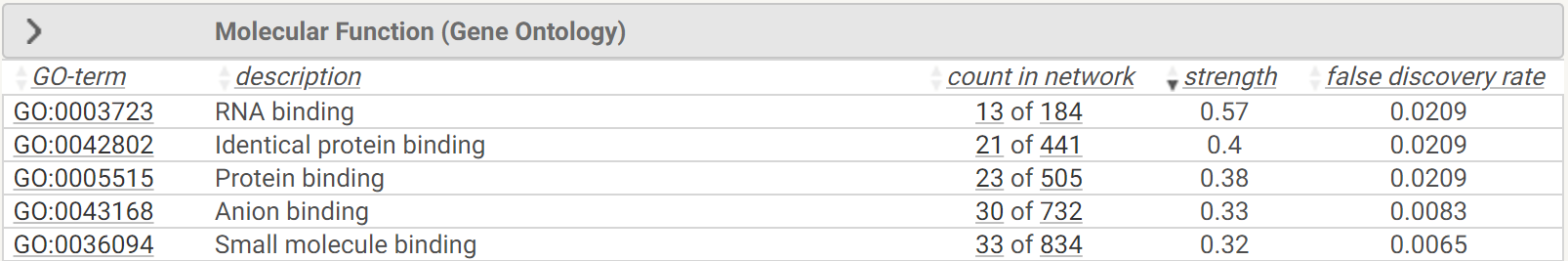

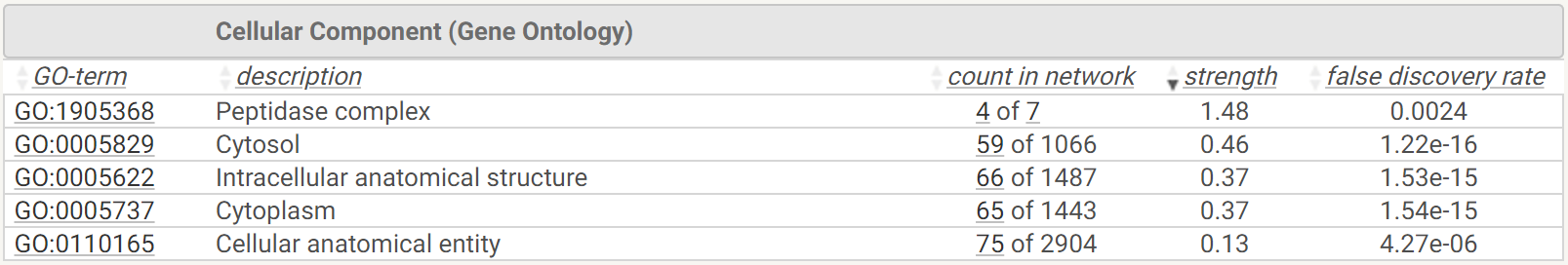

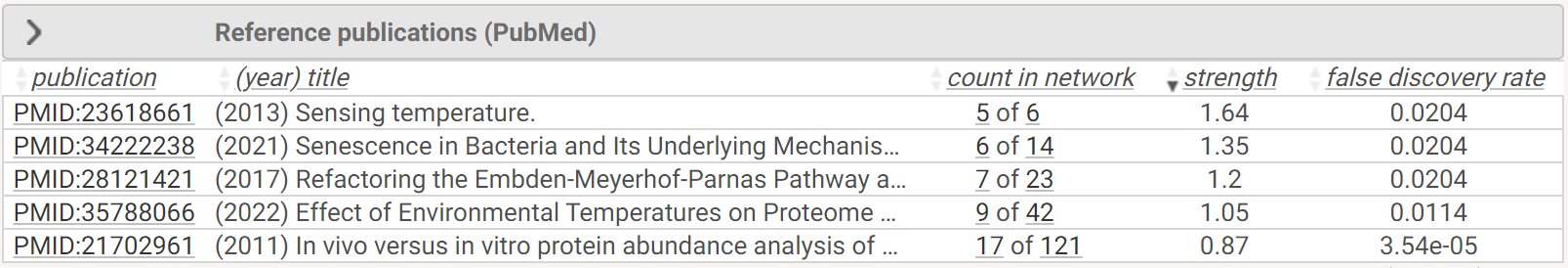

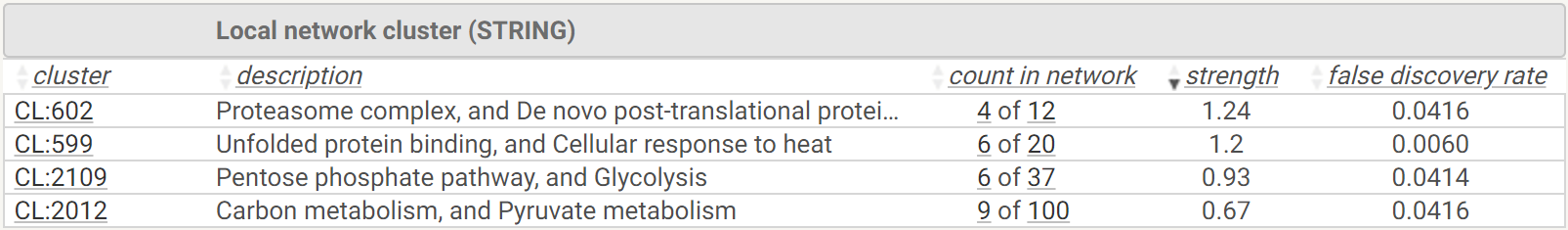

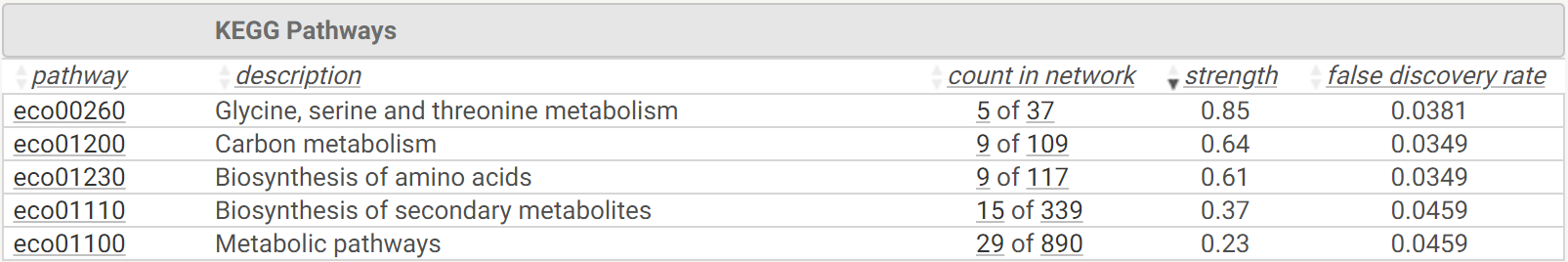

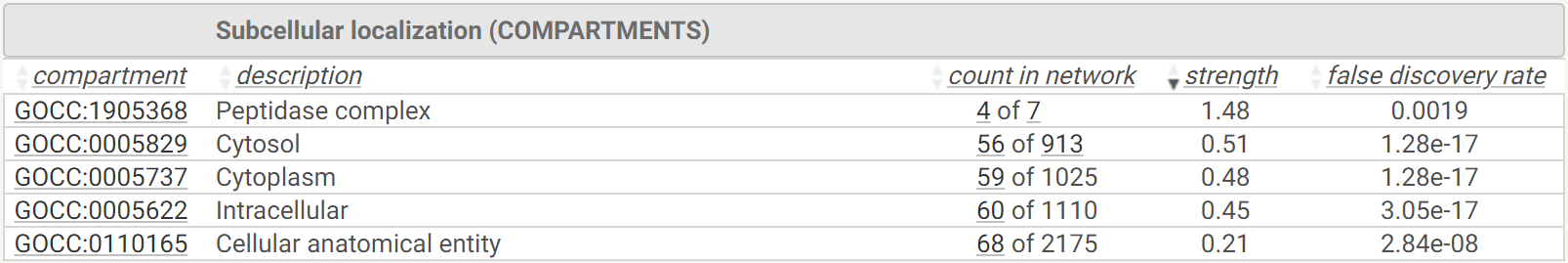

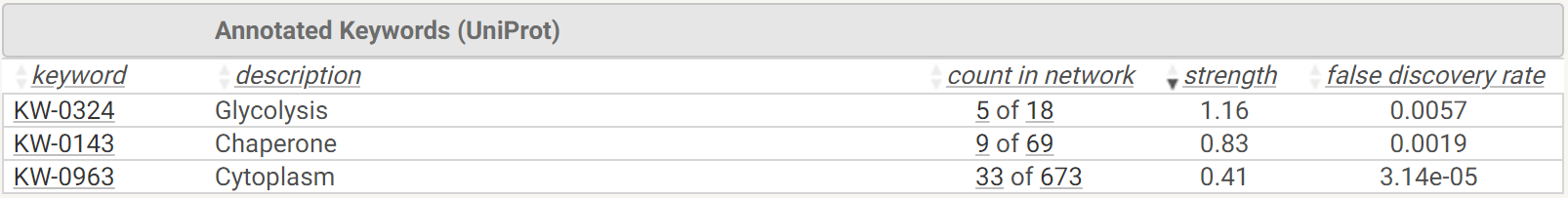

**Table 13.** **The pathway analysis for the downregulated proteins in the Δ*sucA* strain compared to the wild type.** See the legend of Table 9 for further details.

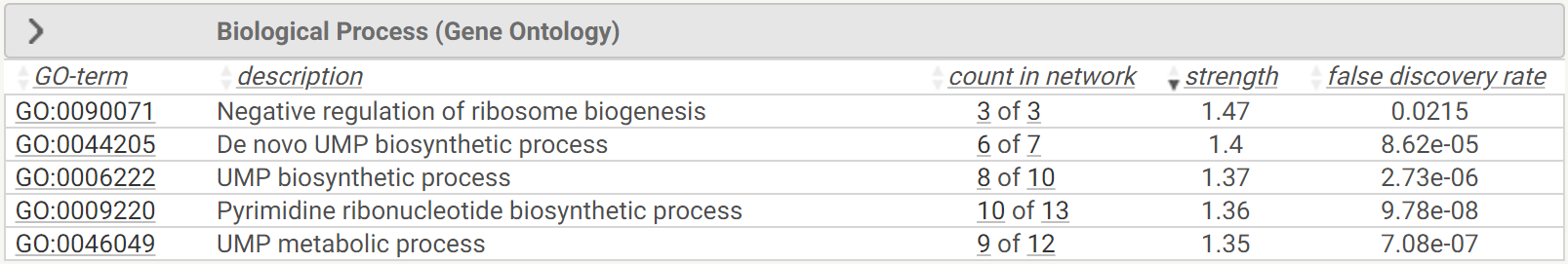

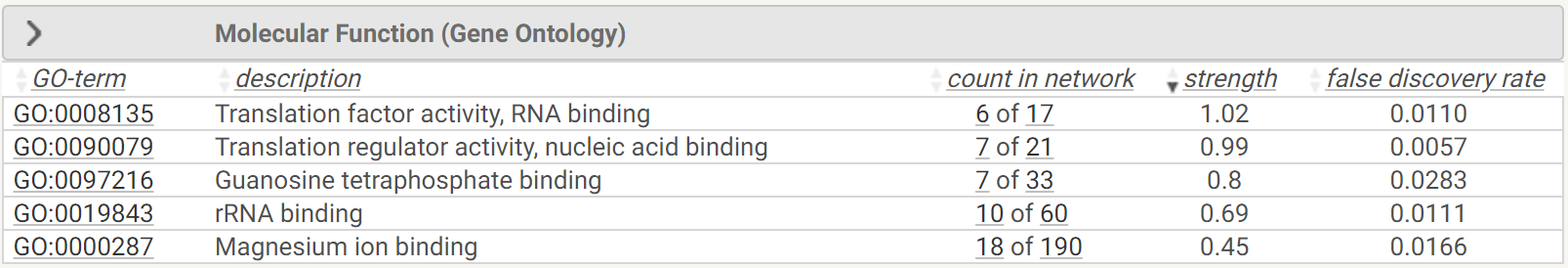

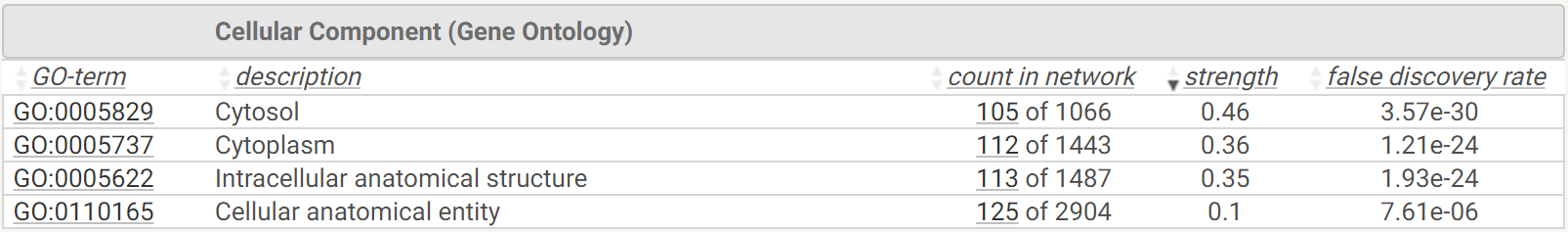

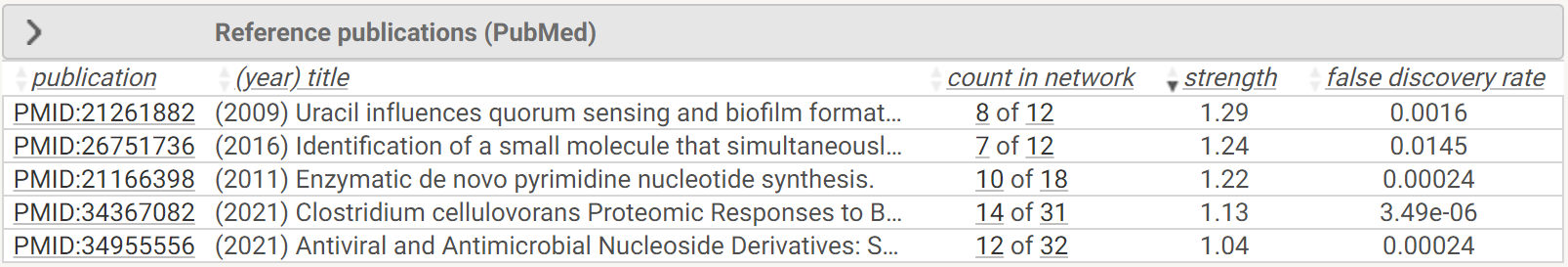

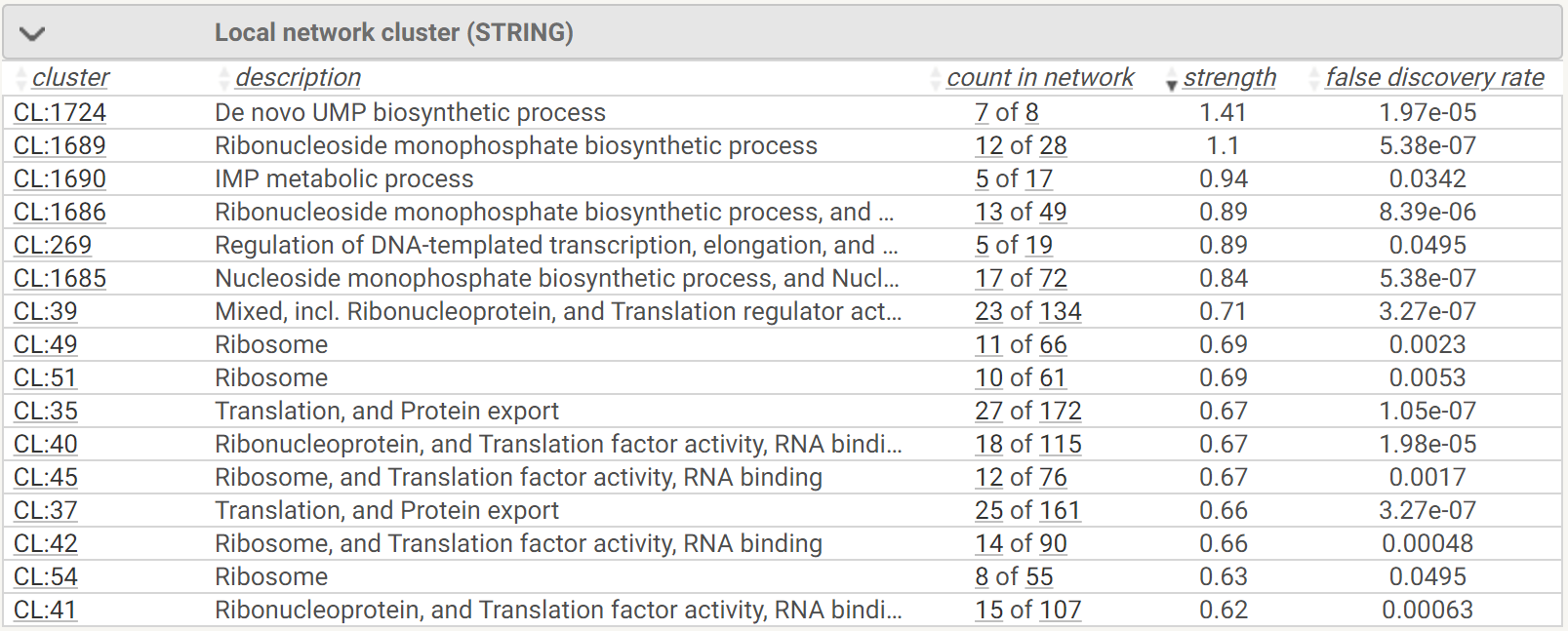

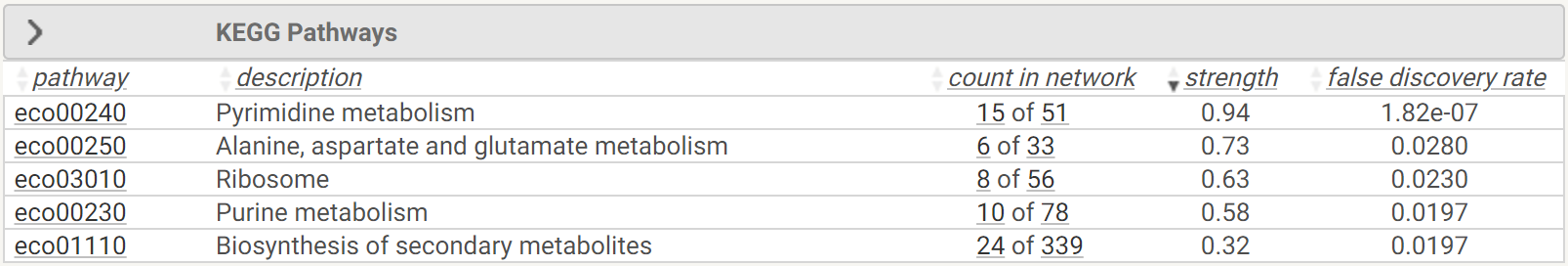

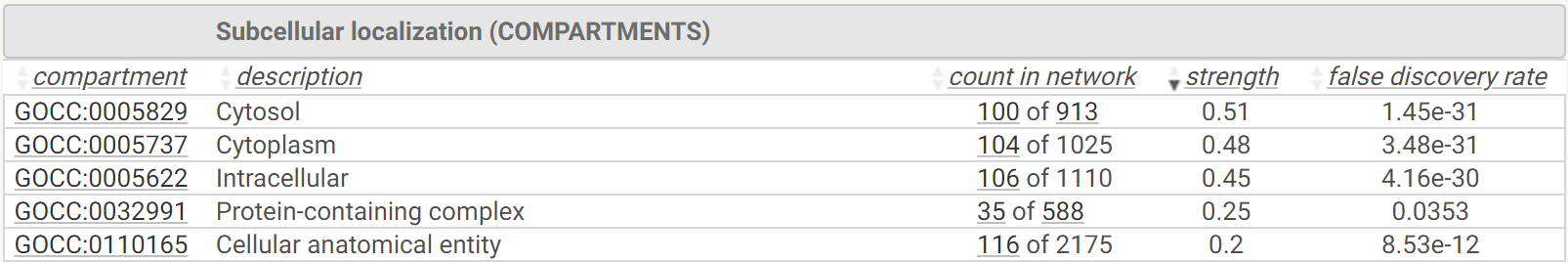

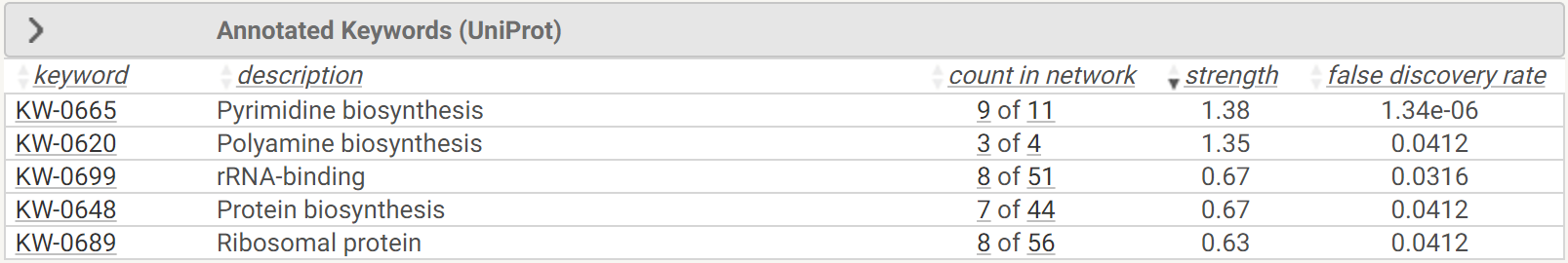

**Table 14.** **The pathway analysis for the downregulated proteins in the Δ*gltA* strain compared to the wild type.** See the legend of Table 9 for further details.

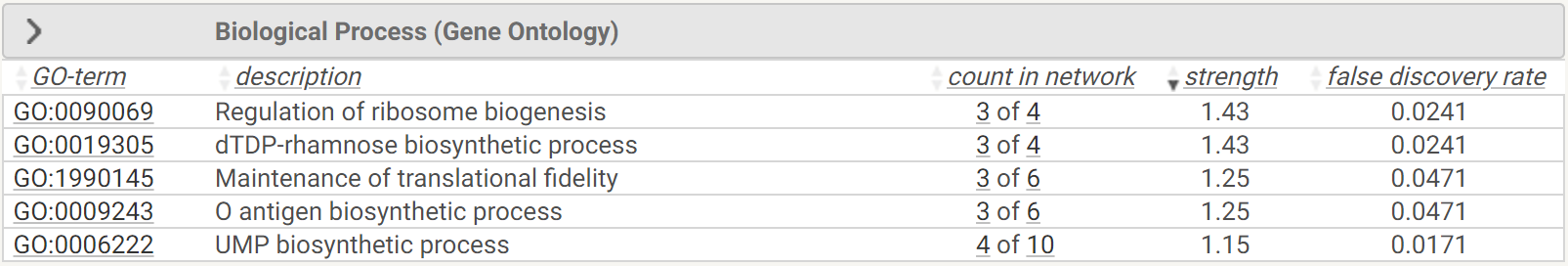

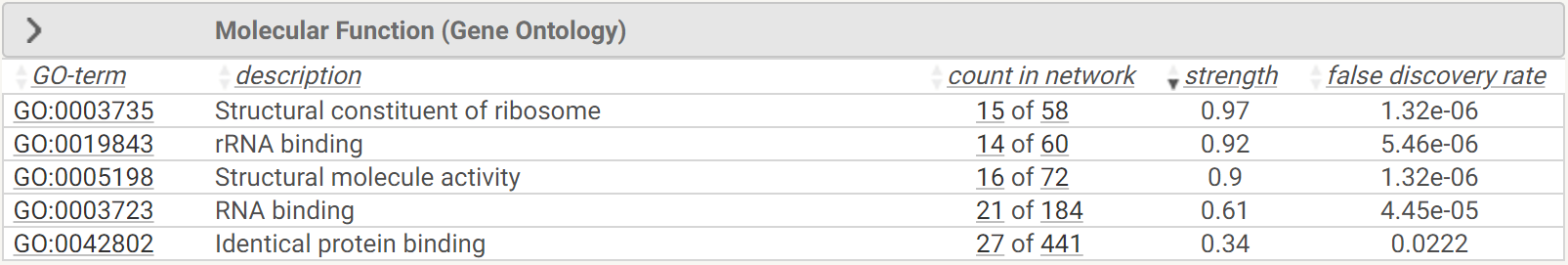

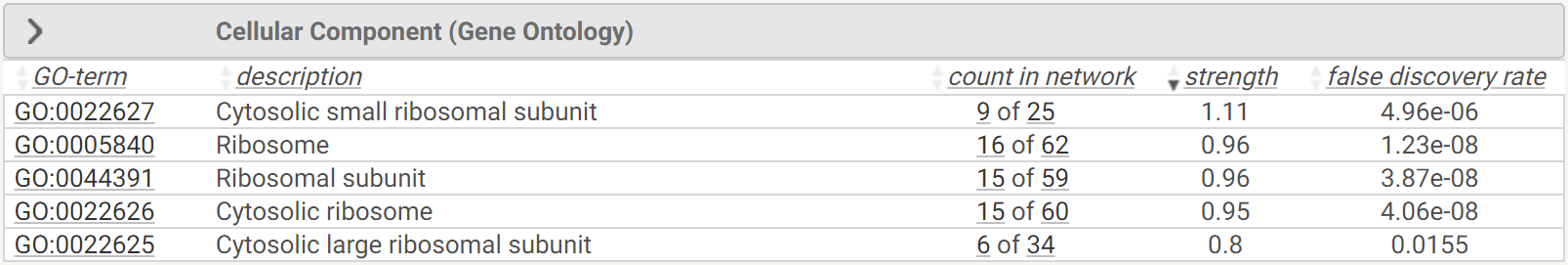

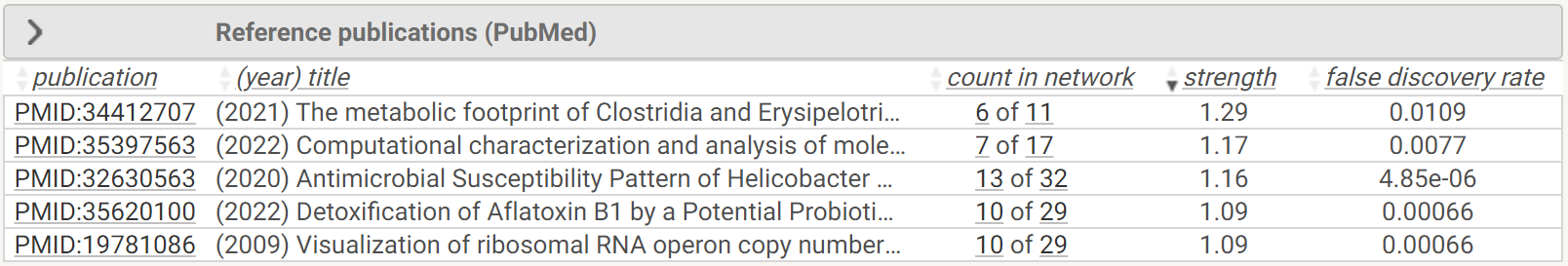

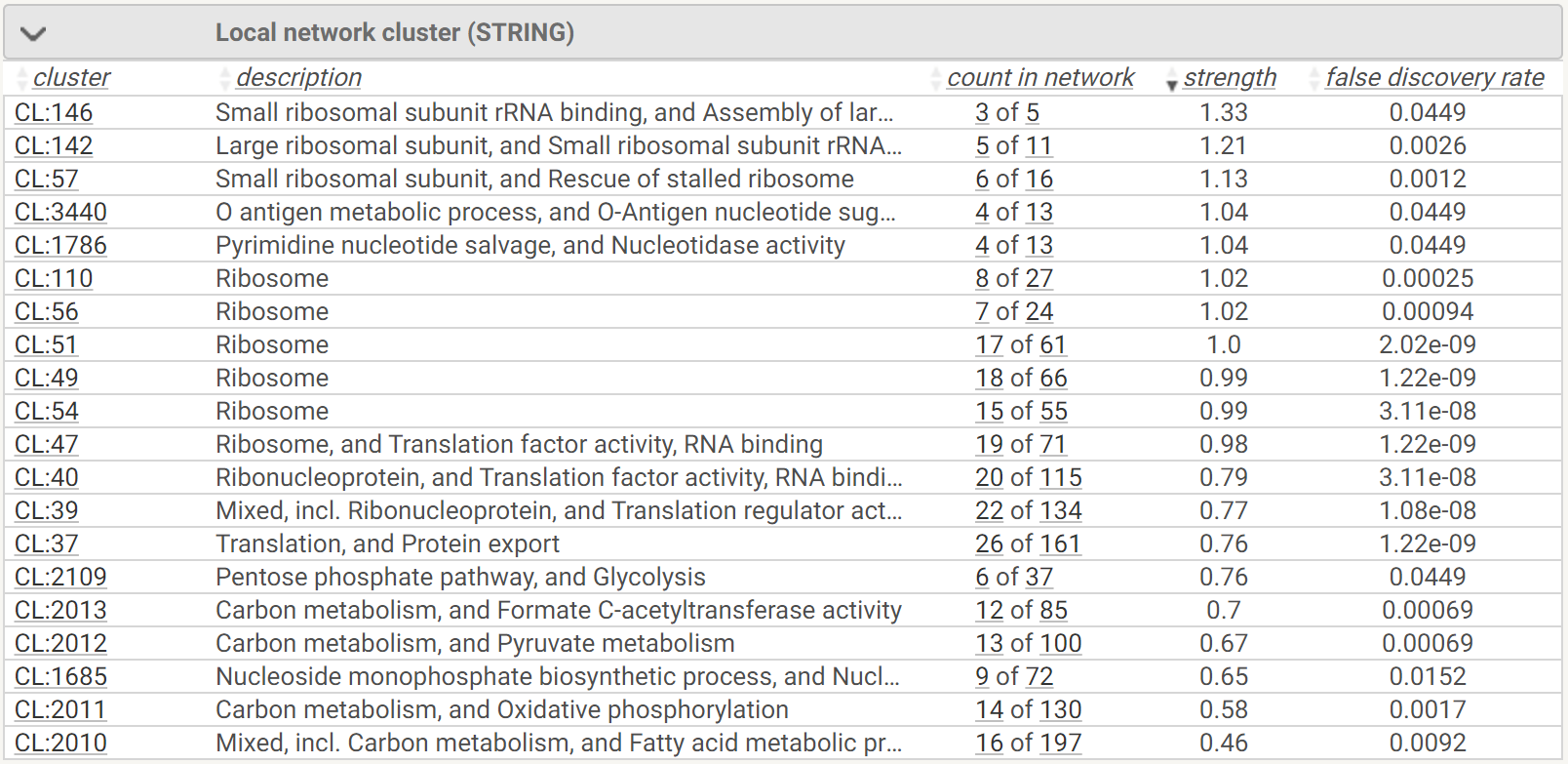

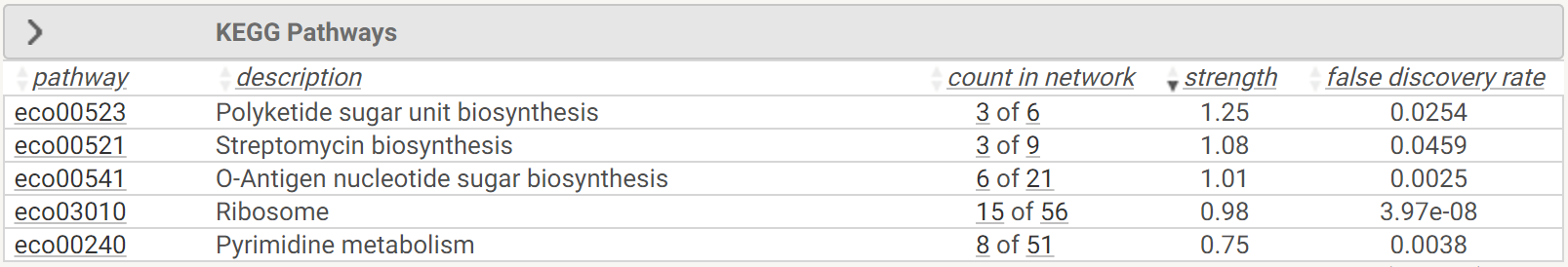

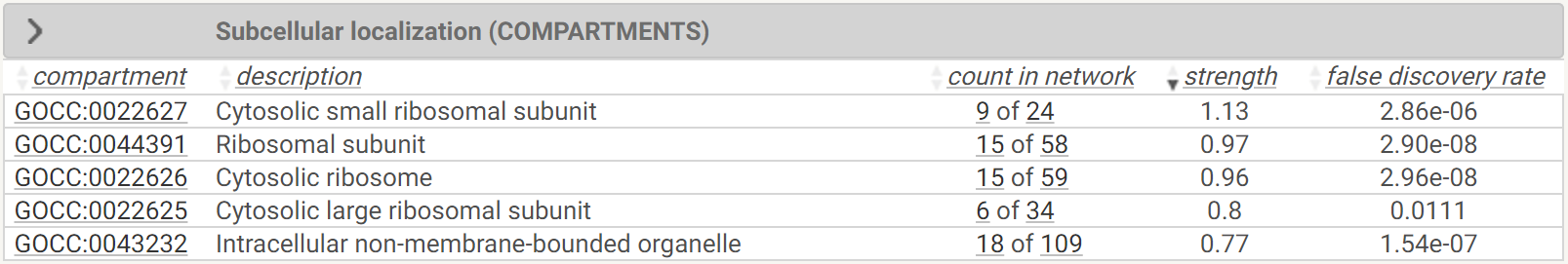

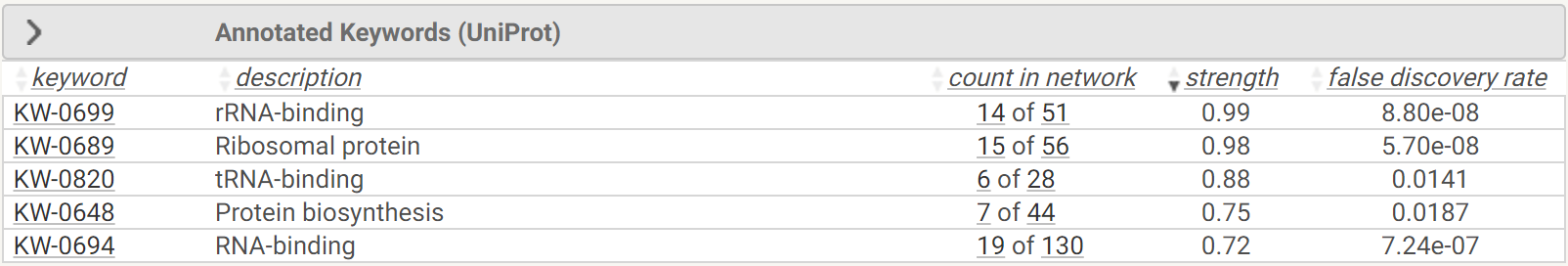

**Table 15.** **The pathway analysis for the downregulated proteins in the Δ*nuoI* strain compared to the wild type.** See the legend of Table 9 for further details.

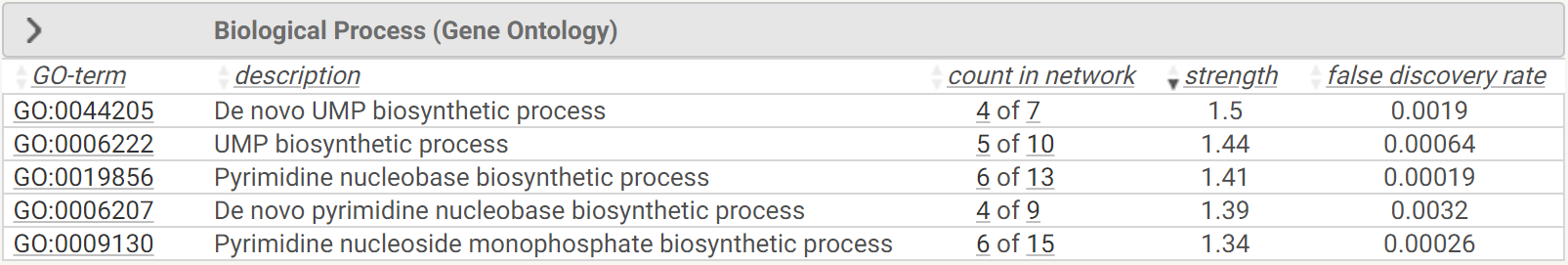

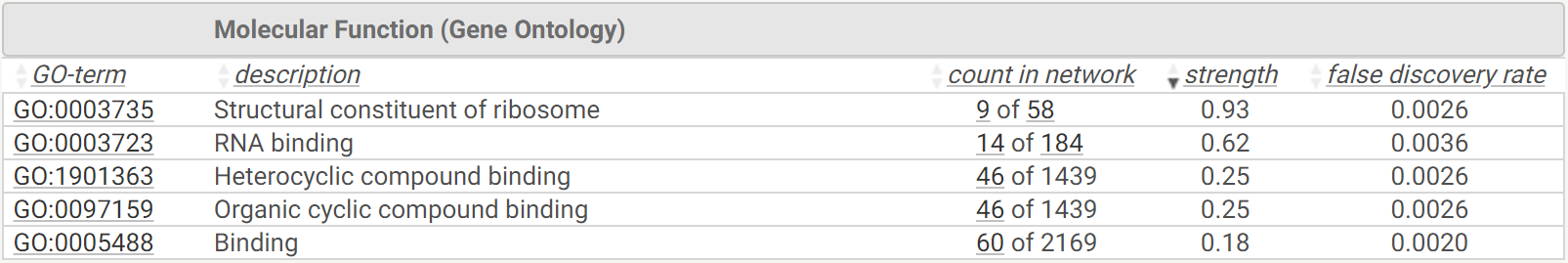

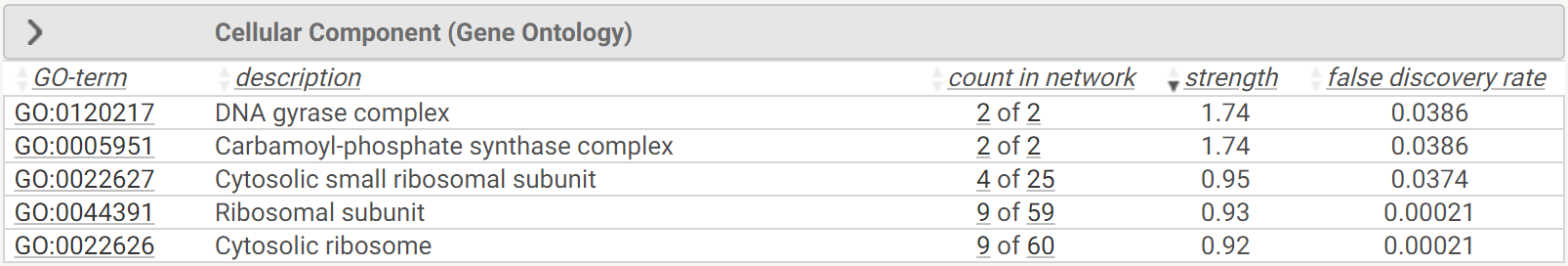

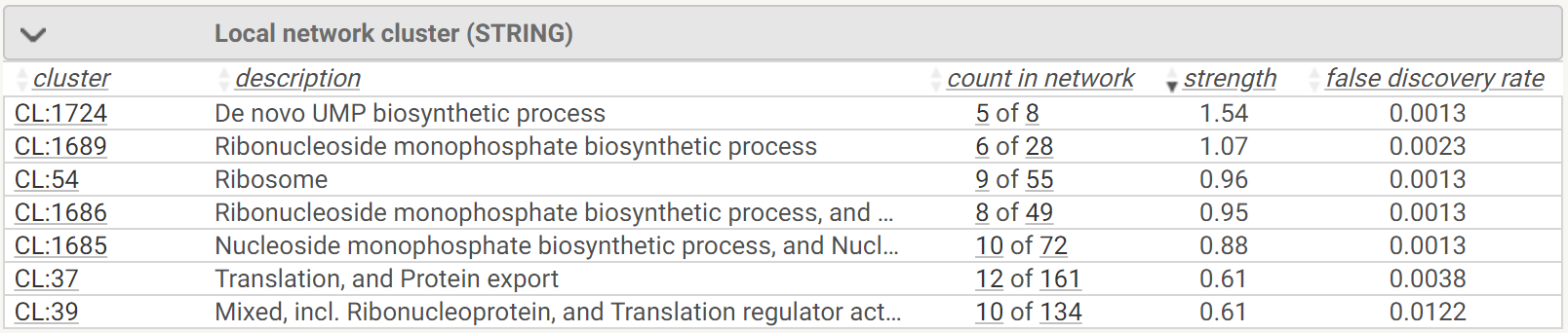

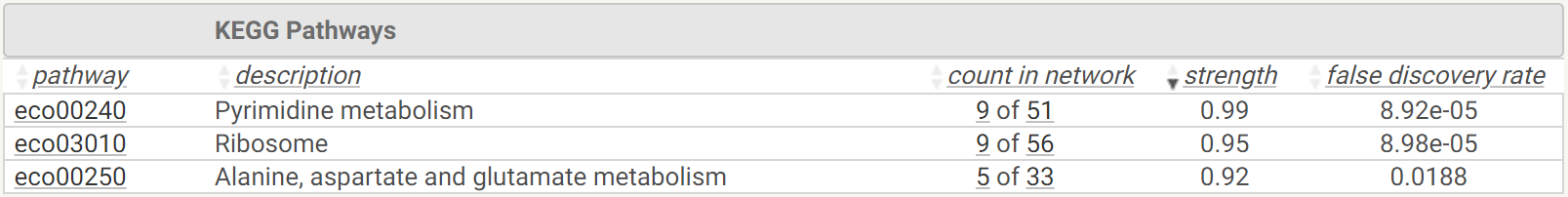

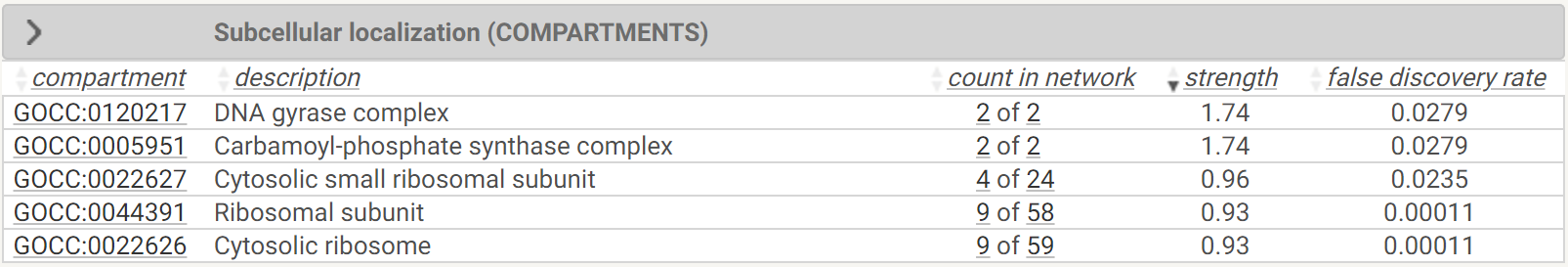

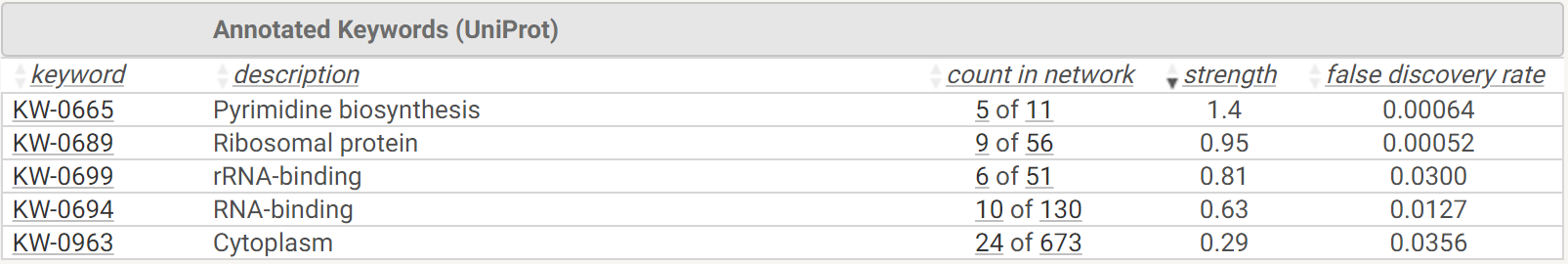

**Table 16.** **The pathway analysis for the downregulated proteins in the Δ*icd* strain compared to the wild type.** See the legend of Table 9 for further details.

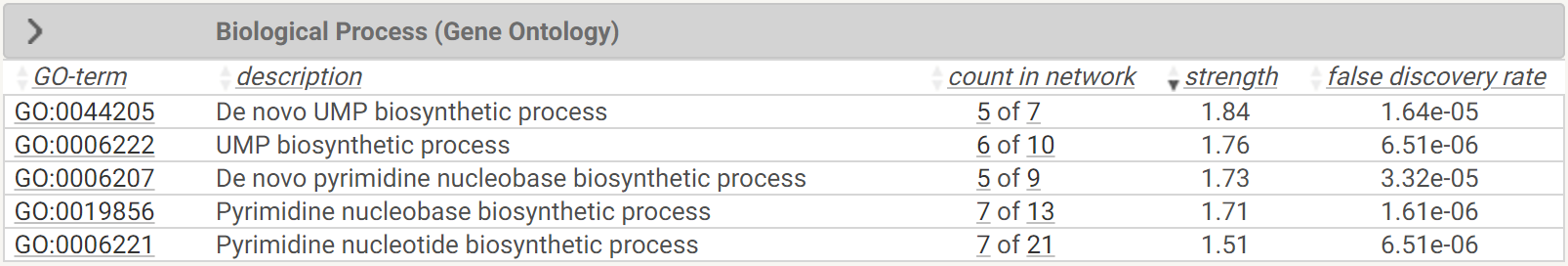

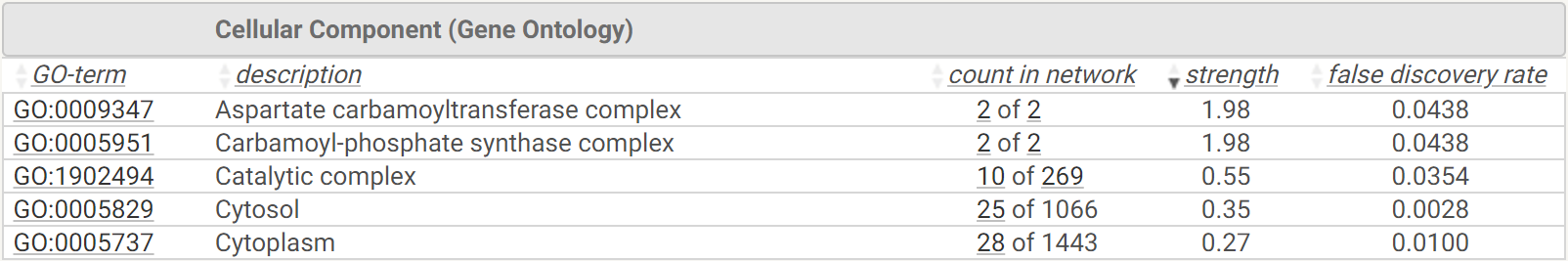

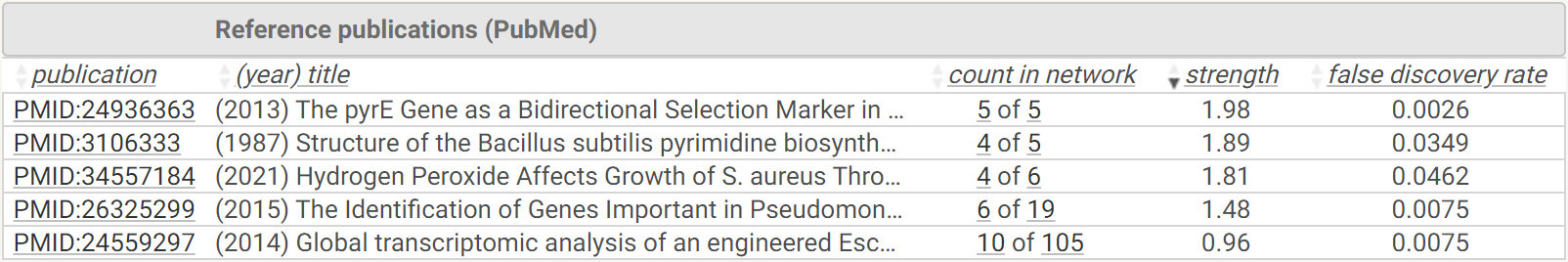

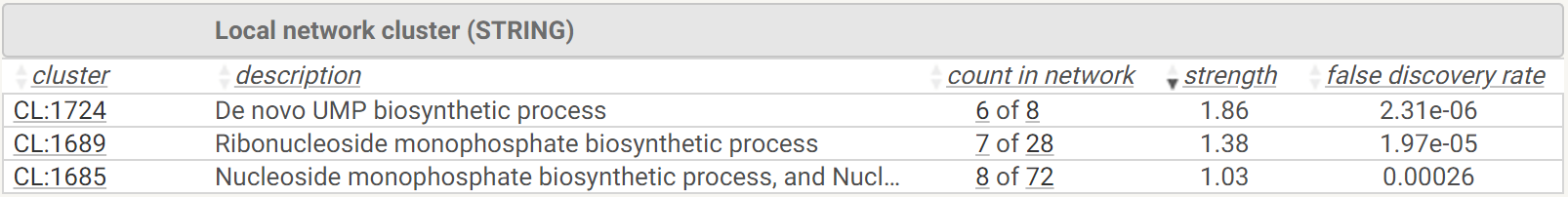

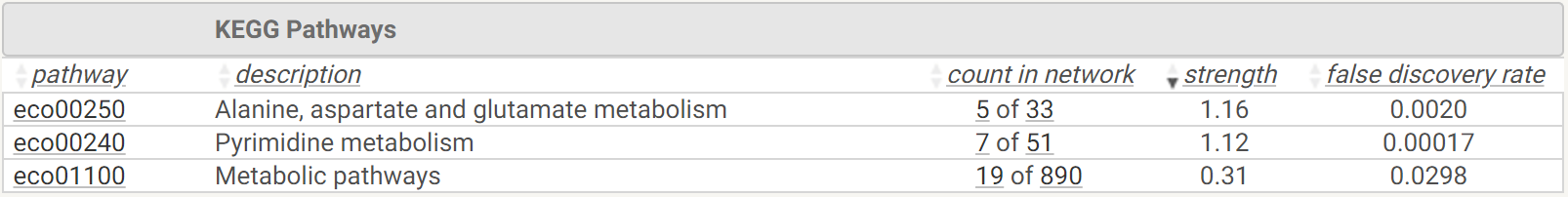

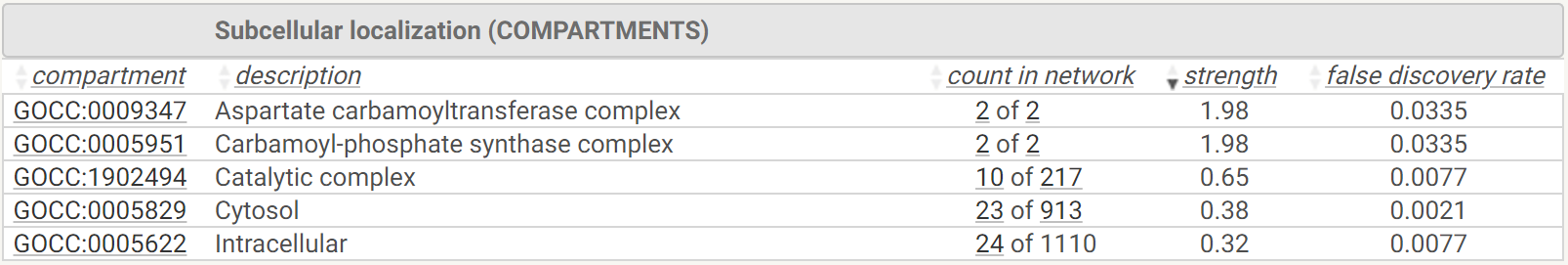

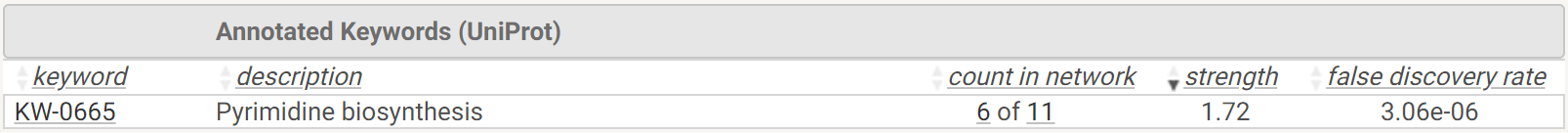
